# Functional characterization of fatty acyl desaturase Fads2 and Elovl5 elongase in the Boddart’s goggle-eyed goby, *Boleophthalmus boddarti* (Gobiidae) suggest an incapacity for long-chain polyunsaturated fatty acid biosynthesis

**DOI:** 10.1101/751057

**Authors:** Han-Jie Soo, Joey Chong, Lau Nyok Sean, Seng Yeat Ting, Sam Ka Kei, Meng-Kiat Kuah, Sim Yee Kwang, M. Janaranjani, Alexander Chong Shu-Chien

## Abstract

Long-chain polyunsaturated fatty acid biosynthesis, a process to convert C18 polyunsaturated fatty acids to eicosapentaenoic acid (EPA), docosahexaenoic acid (DHA) or arachidonic acid (ARA) requires the concerted activities of two enzymes, the fatty acyl desaturase (Fads) and elongase (Elovl). This study highlights the cloning, functional characterisation and tissue expression pattern of a Fads and Elovl from the Boddart’s goggle-eyed goby (*Boleophthalmus boddarti*), a mudskipper species widely distributed in the Indo-Pacific region. Phylogenetic analysis revealed that the cloned Fads and Elovl are clustered with other teleost Fads2 and Elovl5 orthologs, respectively. Interrogation of the genome of several mudskipper species, namely *B. pectinirostris, Periophthalmus schlosseri* and *P. magnuspinnatus* revealed a single Fads2 for each respective species while two elongases, Elovl5 and Elovl4 were detected. Using a heterologous yeast assay, the *B. boddarti* Fads2 was shown to possess low desaturation activity on C18 PUFA. In addition, there was no desaturation of C20 and C22 substrates. In comparison, the Elovl5 showed a wide range of substrate specificity, with capacity to elongate C18, C20 and C22 PUFA substrates. We identified an amino acid residue in the *B. boddarti* Elovl5 that affect the capacity to bind C22 PUFA substrate. Both genes are highly expressed in brain tissue. Among all tissues, DHA is highly concentrated in neuron-rich tissues while EPA is highly deposited in gills. Taken together, the results showed that due to disability of desaturation steps, *B. boddarti* is unable to biosynthesis LC-PUFA, relying on dietary intake to acquire these nutrients.

## Introduction

Long chain polyunsaturated fatty acids (LC-PUFA) which include eicosapentaenoic acid (EPA; 20:5n-3), docosahexaenoic acid (DHA; 22:6n-3) and arachidonic acid (ARA; 20:4n-6) are important for energy, cellular membrane integrity, signaling and transcriptional regulation (Jump, 2002). In humans, sufficient consumption of n-3 LC-PUFA is required to ensure healthy cardiovascular functions, anti-inflammatory activities and control of psychiatric diseases (de Deckere, 2001, Arts et al., 2001). In vertebrates, LC-PUFA can be obtained from dietary intake or conversion of C18 polyunsaturated fatty acids (PUFA). Aquatic organisms are rich sources of EPA and DHA for terrestrial inhabitants due to the presence of primary producers having *de novo* PUFA or in some cases, LC-PUFA biosynthesis activities (Arts et al., 2001, Gladyshev et al., 2009). Subsequent consumers occupying different trophic levels, will further convert PUFA into LC-PUFA. Since aquatic organisms are the main source of LC-PUFA for human population, there is considerable appeal to understand the dietary intake and bioconversion PUFA to LC-PUFA in species occupying different trophic levels (De Troch et al., 2012, Guo et al., 2017).

Capacity for LC-PUFA biosynthesis requires a gamut of fatty acyl desaturases (Fads) and elongases of very long-chain fatty acid (Elovl) enzymes, functioning in a sequential manner to insert a double bond at specific locations of the fatty acyl backbone and to elongate the fatty acyl chain, respectively. Numerous efforts to isolate and functionally characterize these enzymes from a diverse range of fish species have outlined the extent of the diversification of Fads and Elovl (Castro et al., 2016, Garrido et al., 2019). In vertebrates, depending on species, several routes are employed for conversion of linolenic acid (LNA) or linoleic acid (LA) to DHA or ARA, respectively. From LNA to EPA, the Δ6 pathway involves a Δ6 desaturation, followed by elongation and Δ5 desaturation. Another route commence with an elongation step, followed by Δ8 and Δ5 desaturation. From EPA, DHA can be produced using the ‘Sprecher pathway’ where two elongation steps lead to the production of 24:5n-3, followed by a Δ6 desaturation and finally a β-oxidation cleaving (Sprecher et al., 1995). A more direct route involving an elongation step followed by Δ4 desaturation was also unravelled in various taxa groups (Li et al., 2010).

Majority of the characterised Fads and Elovl in vertebrates are from bony fish (Leaver et al., 2008). These work stem from the desire to understand elucidate the consequence of using LC-PUFA-poor vegetable oils as dietary lipid source in aquafeeds. In comparison to most tetrapods which possess *fads1* and *fads2*, Teleostei only possess *fads2* ortholog (Castro et al., 2016). Depending on species, teleostei Fads2 with Δ6, Δ5, Δ8 and Δ4 activities have been reported, in unifunctional, bifunctional or multi-functional orthologs (Hastings et al., 2001, Hastings et al., 2004, Tocher et al., 2006, Kuah et al., 2016). As for elongases, *elovl5* is the principal ortholog isolated from various teleost species and is primarily responsible for the elongation of C18 and C20 PUFA substrates. Another ortholog, the *elovl2* elongase participates in the Sprecher pathway and is limited to small number of species (Morais et al., 2009, Monroig et al., 2009, Oboh et al., 2016, Machado et al., 2018). The diversification of teleost Fads and Elovl is postulated to be the outcome of species adaptation towards the availability of LC-PUFA in their respective natural diet (Morais et al., 2012).

Mudskippers (Order: Perciformes; Family: Gobiidae) are the largest group of amphibious teleost species adapted for terrestrial living through modifications of respiration, ammonia metabolism, vision, immunity and locomotion (You et al., 2014). All mudskippers belong to a monophyletic clade, classified under the subfamily Oxudercinae comprising of 10 genus and 43 species (Murphy and Jaafar, 2017). Within this family, four main genus *Periophthalmus*, *Periophthalmodon*, *Boleophthalmus* and *Scartelaos* represent different levels of adaptations towards terrestrial life (You et al., 2014). *Boleophthalmus boddarti* (Pallas, 1770) or Boddart’s goggle-eyed mudskipper, is an amphibious gobiid mudskipper inhabiting brackish waters of mudflats during high tides (Clayton and Wright, 1989). This species is widely distributed in the Indo-Pacific estuarine regions (Parenti and Jaafar, 2017). While studies on fatty acid composition in sediments, trees, thraustochytrids, molluscs and crustaceans from mangrove ecosystems have been reported, the conversion and transfer of LC-PUFA through different trophic level is still poorly understood (Prosper et al., 2003, Coelho et al., 2011). The elucidation of the capacity of mudskippers for LC-PUFA biosynthesis can potentially provide insights on the importance of LC-PUFA in mudflat environment. In view of the non-existence information on the LC-PUFA biosynthesis capacity in gobies, the isolation and functional characterisation of a Fads and Elov5 from *B. boddarti* are reported here.

## Materials and Methods

### Fish collection and tissue sample preparation

Blue-spotted mudskippers *B. boddarti* were collected from Manjung, Perak, Malaysia (4°10’22”N, 100°39’07”E). Animals were anesthetized with tricaine methanesulfonate (MS-222) prior to dissection. Brain, eye, gill, heart, intestine, liver and skin tissues were dissected for total RNA isolation, kept in RNAlater® solution (Ambion, USA) and stored at −80 °C. The use, handling, maintenance and sacrifice of animals were approved by the USM Institutional Animal Care and Use Committee (USM/IACUC/2019/(117)981).

### RNA isolation and molecular cloning of *B. boddarti* Fads and Elovl full-length cDNAs

Total RNA was isolated from *B. boddarti* liver using TRI Reagent® (Molecular Research Centre, USA) as described in manufacturer’s protocol. Purity and concentration of the isolated RNA were determined using the SmartSpec™ Plus spectrophotometer (BioRad, USA). Integrity of the RNA was examined via 1% (w/v) denaturing formaldehyde agarose gel electrophoresis. Traces of genomic DNA were removed by treating 2 μg of the total RNA with RQ1 RNase-Free DNase (Promega, USA). First-strand cDNA was synthesised from RNA using the M-MLV reverse transcriptase (Promega, USA). Partial cDNAs of the *B. boddarti* desaturase and elongase were obtained using the respective degenerate primers (Table 1). Following this, the cDNA ends of both genes were obtained via 5’- and 3’-RACE using the SMARTer™ RACE cDNA amplification kit (Clontech, USA). Resulting PCR products were directly sequenced and full-length cDNAs of the putative mudskipper desaturase and elongase were constructed by aligning overlapped regions of the cDNA fragments.

**Table 1:**
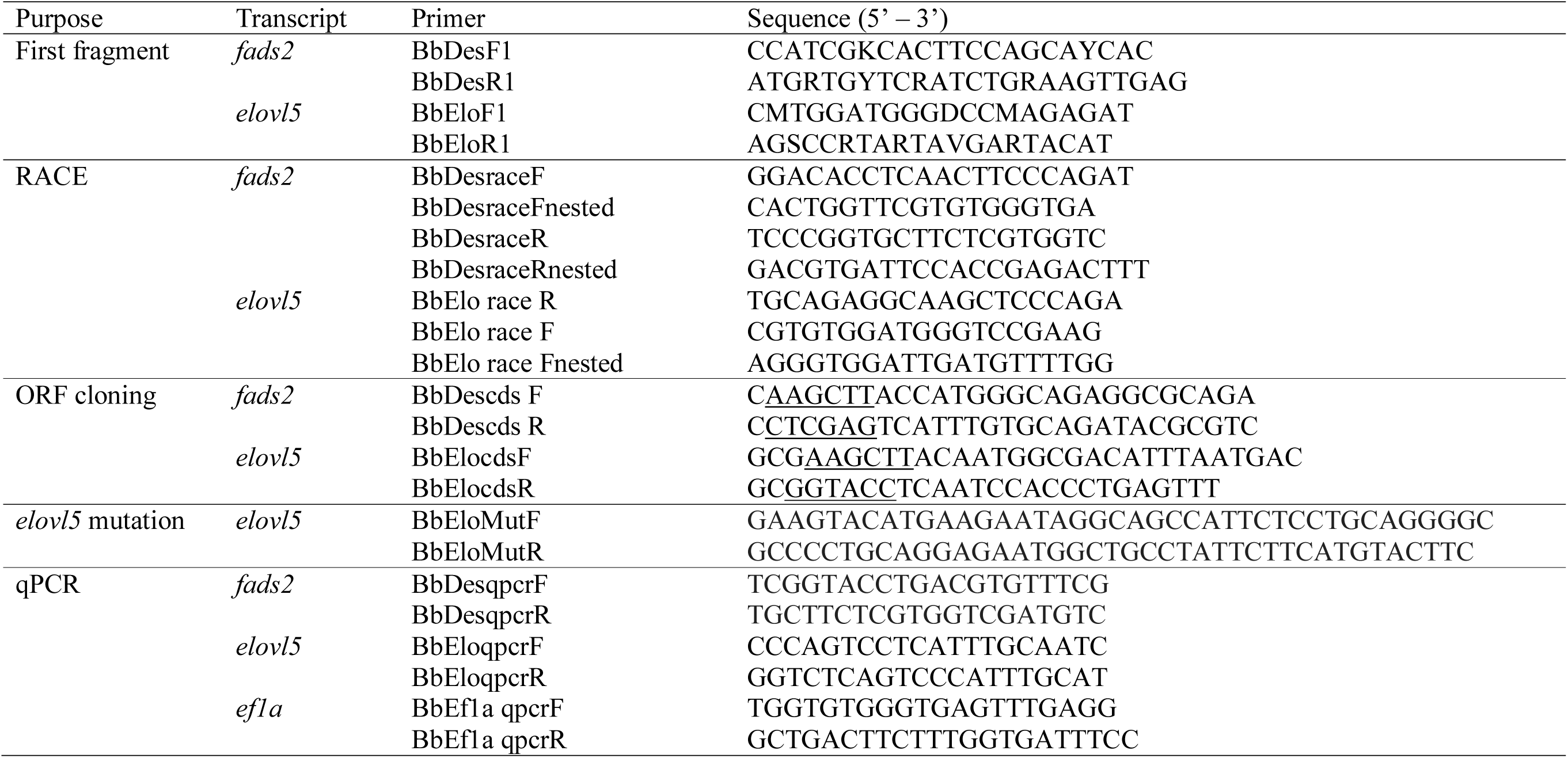
Sequence of primers used in the cloning of *Boleophthalmus boddarti fads2* and *elovl5* ORF, *elovl5* mutation and qPCR tissue expression studies. Restriction sites *HindIII* (AAGCTT), *XhoI* (CTCGAG) and *KpnI*(GGTACC) are shown in underlines.

**Table 2.** In silico mapping of *fads* and *elovl* in genomes of *Boleophthalmus pectinirostris*, *Scartelaos histophorus*, *Periophthalmus magnuspinnatus* and *P. schlosseri*. BLAST similarity was from query against *Danio rerio* and *B. boddarti* orthologs

| Species | Gene | Query | Match in the annotated protein |  |  | Closest match in public database |
| --- | --- | --- | --- | --- | --- | --- |
|  |  |  | Accession no. | Similarity (%) | E-value |  |
| *B. pectinirostris* | *fads2* | XM_020933070.1 | GLEAN_10002267 | 99 | 0.0 | XP_020788729.1 |
|  | *elovl4* | NM_199972.1 | GLEAN_10022389 | 78 | 0.0 | – |
|  | *elovl5* | XM_020921471.1 | GLEAN_10021776 | 100 | 0.0 | XP_020777130.1 |
| *S. histiophorus* | *fads2* | XM_020933070.1 | – | – | – | – |
|  | *elovl4* | NM_200796.1 | GLEAN_10015754 | 80 | 0.0 | – |
|  | *elovl5* | XM_020921471.1 | – | – | – | – |
| *P. schlosseri* | *fads2* | XM_020933070.1 | GLEAN_10005762.1 | 77 | 4e<sup>-99</sup> | – |
|  | *elovl4* | NM_200796.1 | GLEAN_10017052 | 78 | 0.0 | – |
|  | *elovl5* | XM_020921471.1 | GLEAN_10066386 | 91 | 0.0 | – |
| *P. magnuspinnatus* | *fads2* | XM_020933070.1 | GLEAN_10007147 | 80 | 0.0 | ENSPMGT00000023711.1 |
|  | *elovl4* | NM_199972.1 | GLEAN_10010326 | 82 | 1e<sup>-178</sup> | ENSPMGT00000013523 |
|  | *elovl5* | XM_020921471.1 | GLEAN_10017700 | 91 | 0.0 | ENSPMGP00000029011,<br>ENSPMGP00000029023 |

\**B. pectinirostris*: XM\_020933070.1, XM\_020921471.1; *D. rerio*: NM\_199972.1, NM\_200796.1

### Sequence and phylogenetic analysis of *B. boddarti* Fads and Elovl

Obtained sequence of the putative *B. boddarti fads2* and *elovl5* were verified through the BLAST program (http://www.ncbi.nlm.nih.gov/). Multiple sequence alignments of the corresponding amino acid sequences were performed using the Clustal Omega program (http://www.ebi.ac.uk/Tools/msa/clustalo/). The deduced amino acid sequences of both proteins were characterised with the InterProScan program (http://www.ebi.ac.uk/Tools/pfa/iprscan/) and the TOPCONS program (http://topcons.cbr.su.se/).

Mudskipper sequences and orthologs from various species were aligned with MAFFT v7.407 (Katoh and Standley, 2013). Fads sequences of *Gnathonemus petersii, Megalops cyprinoides* and *Osteoglossum bicirrhosum* were kindly provided by Lopes-Marques, M. (personal communication). The software returned the L-INS-I as the probably most accurate alignment method (Katoh et al., 2005). Alignment was subjected to smart model selection (SMS) and Maximum Likelihood (ML) phylogenetic evolutionary tree was constructed with PhyML 3.0 using Abayes as support (Lefort et al., 2017, Anisimova et al., 2011, Guindon et al., 2010). These analysis were conducted using the NGPhylogeny.fr integrative web service (Lemoine et al., 2019). The resulting tree was visualised using Fig-Tree v1.3.1 (http://tree.bio.ed.ac.uk/software/figtree/).

### Functional characterisation of *B. boddarti* Fads and Elovl through heterologous expression in yeast

ORFs of the cloned *fads* and *elovl* were amplified from their respective first-strand cDNA using a forward primer containing a restriction site for *Hind* III and a reverse primer containing a restriction site for *Xho* I (Table 1). PCR products were cloned into the pGEM^®^-T Easy vector (Promega, USA). DNA fragments containing ORFs were digested with restriction enzymes (New England BioLabs, UK) and ligated into restricted pYES2 yeast expression vectors (Invitrogen, USA). Ligated products were transformed into *E. coli* DH5α competent cells, followed by screening of colonies for successful insertion. After purification, pYES2 vector containing the ORF was transformed into *Saccharomyces cerevisiae* yeast strain INVSc1 competent cells using the S.c. EasyComp™ Transformation kit (Invitrogen, USA). Screening for pYES2 construct-transformed yeast colonies was carried out using the *S. cerevisiae* minimal medium without uracil (SCMM-U) and confirmed using PCR.

For heterologous functional characterisation study, a single colony was selected from the desired recombinant yeast strain and propagated overnight in SCMM-U broth containing 2% (v/v) raffinose at 30°C at 250 rpm. Subsequently, the overnight yeast culture was diluted to an OD_600_ of 0.4 in new SCMM-U broth and allowed to grow until the OD_600_ reached 1. At this point, galactose was added to the culture medium at a concentration of 2% (v/v) for induction of expression. In order to functional characterise the cloned ORF, the yeast culture medium was supplemented with specific fatty acid substrate: 0.5 mM for C18, 0.75 mM for C20 and 1.0 mM for C22. For *B. boddarti* Fads2, ALA, LA, 20:3n-3, 20:4n-3, 20:2n-6, 20:3n-6, 22:5n-3 and 22:4n-6 were evaluated while for Elovl5, ALA, LA, 18:4n-3, 18:3n-6, EPA, ARA, 22:5n-3 and 22:4n-6 were tested. After 48 hours of incubation, yeast cells were harvested via centrifugation of 5 min at 4,000 × *g*. The resulted cell pellet was washed twice and stored at −80°C before lipid extraction.

### P57Q Mutation of *B. boddarti* Elovl5

Mutation of the cloned *B*. *boddarti* Elovl5 sequence was carried out using the QuikChange site-directed mutagenesis kit (Agilent Technologies, La Jolla, CA) according to the manufacturer’s protocol with primers listed in Table 1 using the pGEM®-T Easy vector containing the *B. boddarti elovl5* sequence as template (Table 1). Plasmid extracted from the positive clone was digested and ligated into pYES2 vector. The sequence of this P57Q construct was confirmed by DNA sequencing prior to transformation into *S. cerevisiae*. In vitro assay to determine the functional capacity of the mutated *B. boddarti* was carried out as described above.

### Total lipid extraction and FAME analysis via GC-MS

Total lipids were extracted from harvested yeast cells via sonication in a mixture of chloroform and methanol at ratio of 2:1 (v/v), followed by methylation and transesterification using boron trifluoride in methanol. The resulting fatty acid methyl esters (FAME) were analysed and quantified using a GC-MS QP2010 Ultra gas chromatograph-mass spectrometer (Shimadzu, USA), equipped with a BPX70 high-polarity fused-silica capillary column (60 m length, 0.25 mm inner diameter, 0.25 μm film thickness; SGE, USA). Helium was supplied as carrier gas and oven temperature was programmed from 100 °C to 210 °C at a rate of 2 °C/min and held at 210 °C for 30 min. Temperatures of the injector and detector were set at 250 and 230 °C, respectively. Individual FAME was identified through mass spectrometry based on the NIST08s mass library (Shimadzu, USA) and GC retention time. Desaturation or elongation conversion efficiencies of respective added fatty acid were calculated as the proportion of exogenously added fatty acid to the desaturated or elongated FA products, [product area/(product area + substrate area)] × 100. For every substrate, the assay was repeated thrice with different recombinant yeast isolate.

### Tissue expression *B. boddarti fads2* and *elovl5* via qPCR analysis

Total RNA was isolated from adult mudskipper tissues using TRI Reagent® (Molecular Research Centre, USA), followed by measurement of concentration, purity and integrity. Treatment with RQ1 RNase-Free DNase (Promega, USA) was carried out, followed by qPCR analysis on a 7500 Real-Time PCR System with the Power SYBR® Green RNA-to-C_T_™ 1-Step kit (Applied Biosystems, USA) using primers listed in Table 1. Gene-specific standard curves were generated from a dilution series constructed from RNA pooled from all experimental samples. Specificity of PCR amplification was examined through melting curve analysis via continuous fluorescence acquisition from 60°C to 95°C at a transition rate of 0.05 °C/s. An amplification reaction without RNA as template was used as negative control. Relative expression level of target genes was computed using the Gene Expression Macro™ version 1.1 (BioRad, USA) with the average CT value normalised to the internal control gene *ef1a* (ALE 14778). The qPCR data are expressed as the mean ± SEM from three individual adult fish. Statistical comparison of expression levels between different tissues was conducted using the one-way ANOVA followed by a Tukey’s post hoc test (P < 0.05).

### Fatty acid composition of *B. boddarti* tissues

Liver, intestine, muscle, eye, gill, gonad, heart and brain tissues were dissected from adult mudskippers and kept at −80 °C. Total lipids were extracted from tissues (0.5-1.0 g per replicate) followed by analysis for fatty acid composition as described earlier.

### In silico search for *fads* and *elovl* genes in *B. boddarti* genome

The annotated gene models for *B*. *pectinirostris* (blue-spotted mudskipper)*, Scartelaos histiophorus* (blue mudskipper), *Periophthalmus schlosseri* (giant mudskipper) and *P*. *magnuspinnatus* (giant-fin mudkipper) were obtained from their published genomes through personal communication (You et al., 2014). Similarity searches on these genomes using blastn were performed with threshold e-value < 1e-10, similarity and coverage at > 70%, respectively. The cloned *B. boddarti* Fads2 and Elovl5 sequences were used as queries. Additionally, *Danio rerio fads*2 (NP 571720.2) and Elovl 2 (AAI29269.1), Elovl4 (NP 957090) and Elovl5 (NP 956747.1) sequences were also used as queries.

## Results

### Cloning, sequence and phylogenetic analyses of *B. boddarti Fads2* and *Elovl5*

Full-length cDNAs of putative *B. boddarti* desaturase and elongase were isolated from the liver tissue of mudskipper. The full-length cDNA of the *fads2* desaturase is 1590 bp, contained an ORF of 1311 bp encoding a putative protein of 436 aa and was deposited in the GenBank database (ALE14476.1) The deduced amino acid sequence of this Fads2 contained all the characteristic features of a microsomal fatty acyl front-end desaturase, such as a N-terminal cytochrome-b5 domain with a heme-binding motif HPGG, three histidine-rich boxes and four transmembrane regions (Fig. 1). In addition, a four amino acid residues FHLQ corresponding to PUFA substrate selectivity was identified within the third transmembrane region. The position of the cloned *B. boddarti* Fads on the phylogenetic tree revealed a strong grouping with Fads2 from other Teleostei representatives (Fig. 2). The clade with the four mudskipper (Gobiformes) Fads2 sequences shares a same ancestor with a large clade of percomorphans, which includes representatives from Cichliformes, Perciformes, Atheriniformes, Beloniformes and Pleuronectiformes. This Percomorpha clade is supported by a cod (*Gadus morhua*) Fads2 sequence. Species from the order Salmoniformes and Osteoglossiformes also formed their respective clades. There was also a strong cluster of the ancient lineage of Clupeocephala represented by several cyprinids, a Siluriformes and a Characiformes, respectively. The tree also showed a clear separation between the vertebrate Fads2 and Fads1, with the later represented by several mammals, and several early Teleostei species.

**Fig 1.**
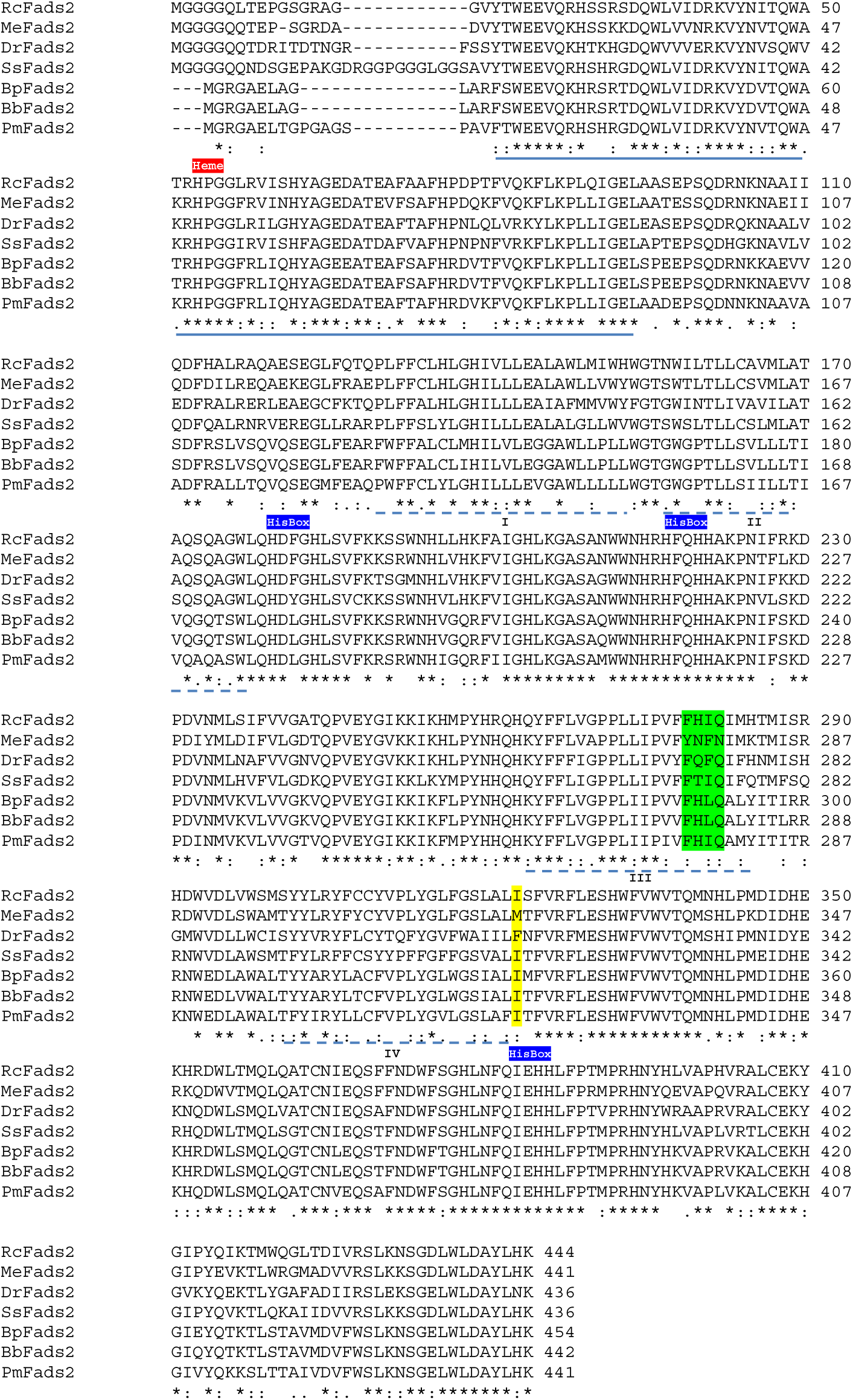
Comparison of Fads2 amino acid sequences from *Boleophthalmus boddarti* (Bb, ALE14476.1) *B. pectinirostris* (Bp, XP020788729.1), *Periophthalmus magnuspinnatus* (Pm, ENSPMGT00000023711.1), *Danio rerio* (Dr, AAG25710.1), *Rachycentron canadum* (Rc, ACJ65149.1), *Salmo salar* (Ss, AAR21624.1) and *Menidia estor* (Me, AHX39206.1). Identical, strongly and weakly similar residues are marked with asterisks, colons and full stops, respectively. The cytochrome-b5 domain is underlined with a solid line while the four putative transmembrane regions are underlined with dotted lines. The three histidine boxes and the heme-binding motif are labeled. Residues corresponding to substrates specificity and regioselectivity are highlighted in green and yellow.

**Fig 2.**
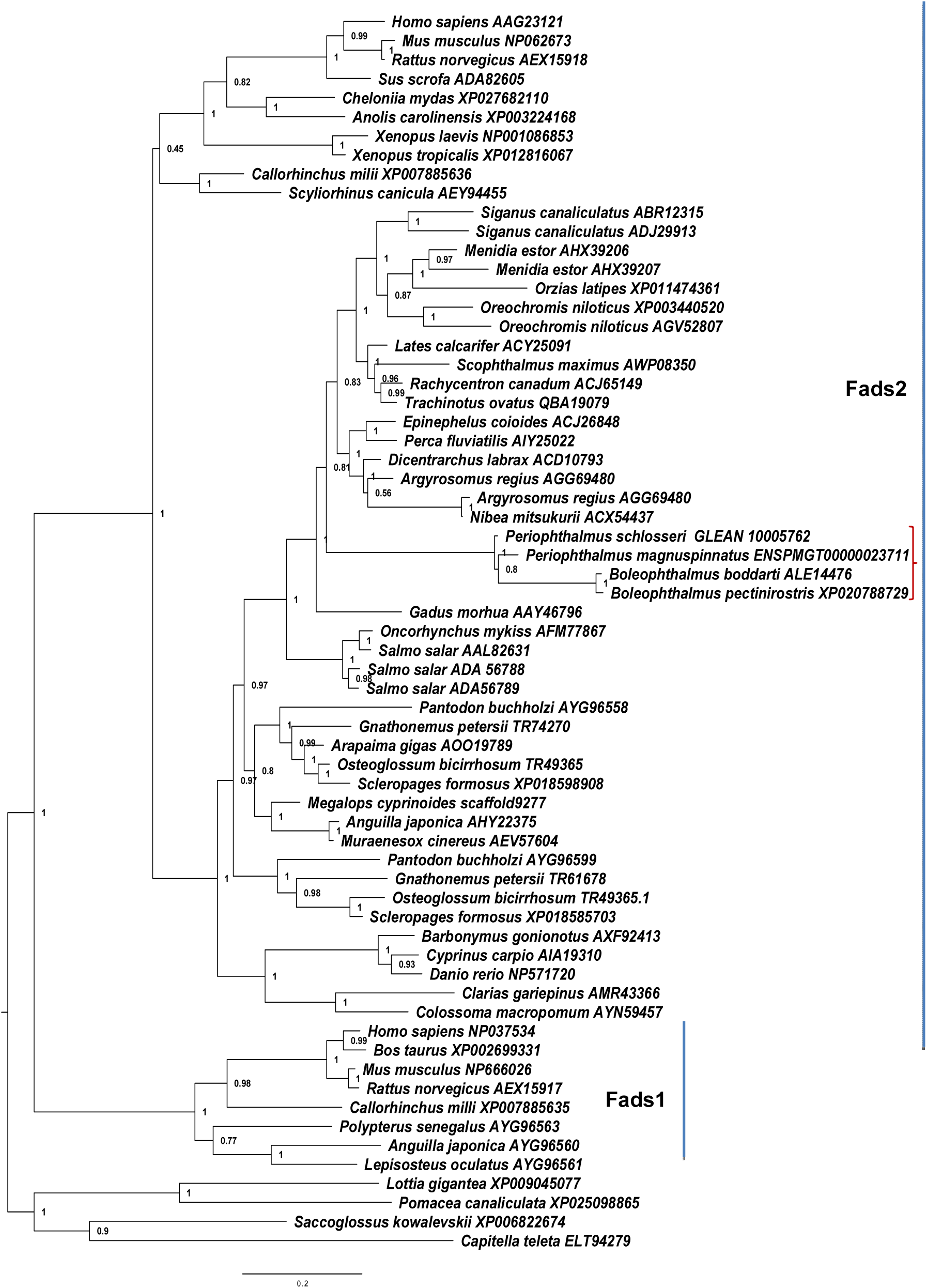
Maximum Likelihood Phylogenetic analysis, using Abayes as support, of 65 Fads desaturase amino acid sequences from various vertebrate species. The tree was visualised using Fig-Tree v1.3.1 (http://tree.bio.ed.ac.uk/software/figtree/) and rooted with invertebrate sequences. The four Fads2 sequences from different mudskipper species are indicated. These analysis were conducted using the NGPhylogeny.fr integrative web service.

The cDNA of the *B. boddarti elovl5* elongase was 1387 bp long, containing an ORF of 873 bp encoding protein with 290 amino acid residues (ALE14477.1). Features distinctive to a microsomal fatty acyl elongase were present, including seven transmembrane regions, four conserved motifs (KxxExxDT, QxxFLHxYHH, NxxxHxxMYxYY and TxxQxxQ), a histidine box (HxxHH) and endoplasmic reticulum retention signal residues (Fig. 3). The phylogenetic tree showed distinctive clades of Elovl5, Elovl2 and Elovl4, respectively. The mudskipper elongases are in the Elovl5 and Elovl4 clades (Fig. 4). The cloned *B. boddarti* Elovl belongs to the distinctive Elovl5 clade. Within this clade are two distinct clusters, one which includes tetrapod, cartilaginous fish and a sarcopterygian while another is well represented by the clupeocephalan taxa group. *B. boddarti elovl* is clustered within a large clade which consists of several other percomorphs. In the Elovl4 clade, elongases from *P. magnuspinnatus* and *P. schlosseri* were separated into two distinct branches, respectively.

**Fig. 3:**
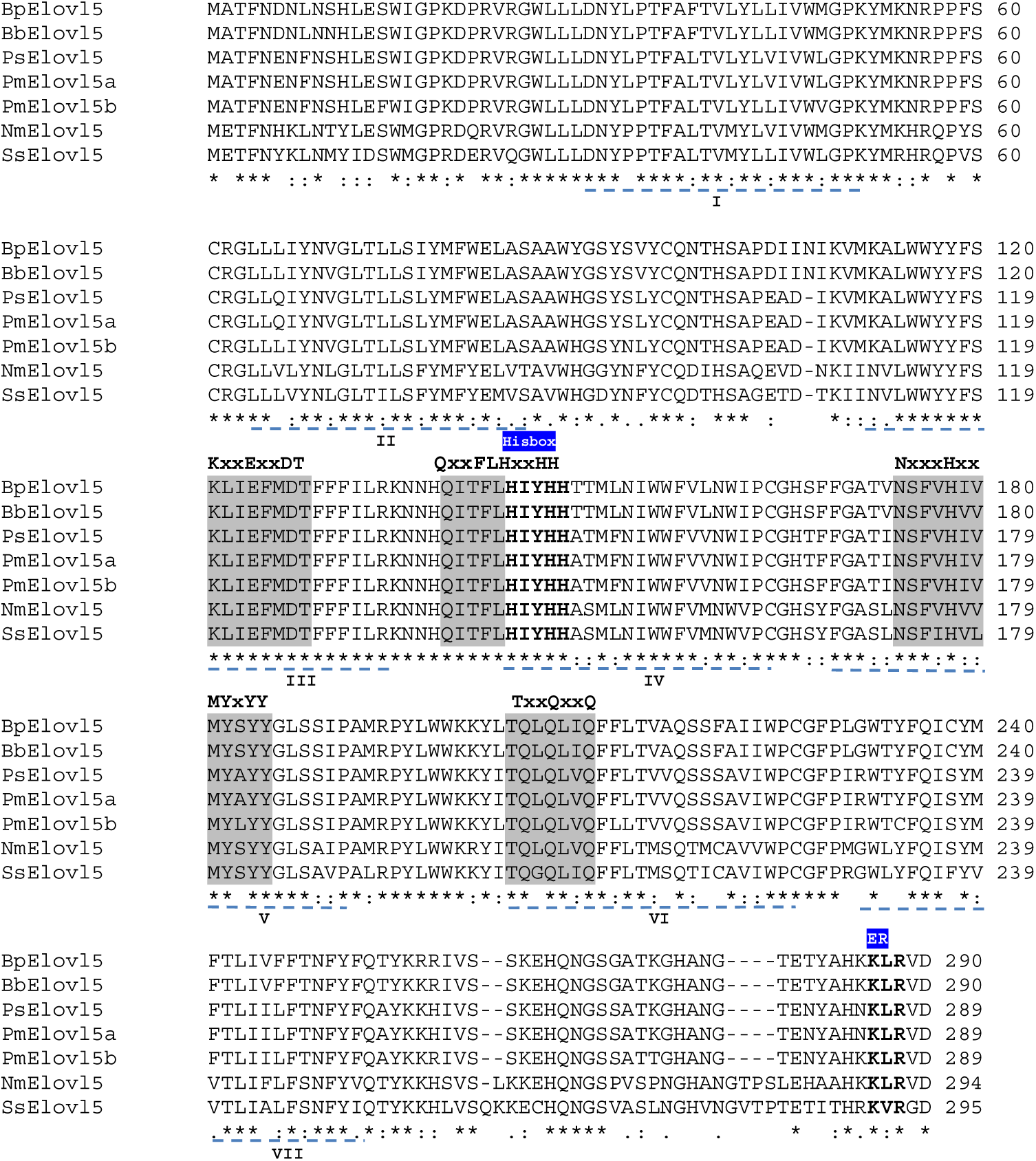
Comparison of the Elovl5 amino acid sequences of *Boleophthalmus boddarti* (Bb, ALE14477.1), *B. pectinirostris* (Bp, XP_020777130.1), *Periopthalmodon schlosseri* (Ps, GLEAN_10066386), *P. magnuspinnatus*, (Pm ENSPMGP00000029011, ENSPMGP00000029023) *Nibea mitsukurii* (Nm, ACR47973.1) and *Salmo salar* (Ss, NP_00111739.1). Identical residues are marked by asterisks, whereas strongly and weakly similar residues are marked by colons and full stops, respectively. The five putative transmembrane regions are underlined with dotted lines. The endoplasmic reticulum retention signal (ER) and the histidine box (HxxHH) are shown. The four conserved motifs are highlighted in grey.

**Fig 4.**
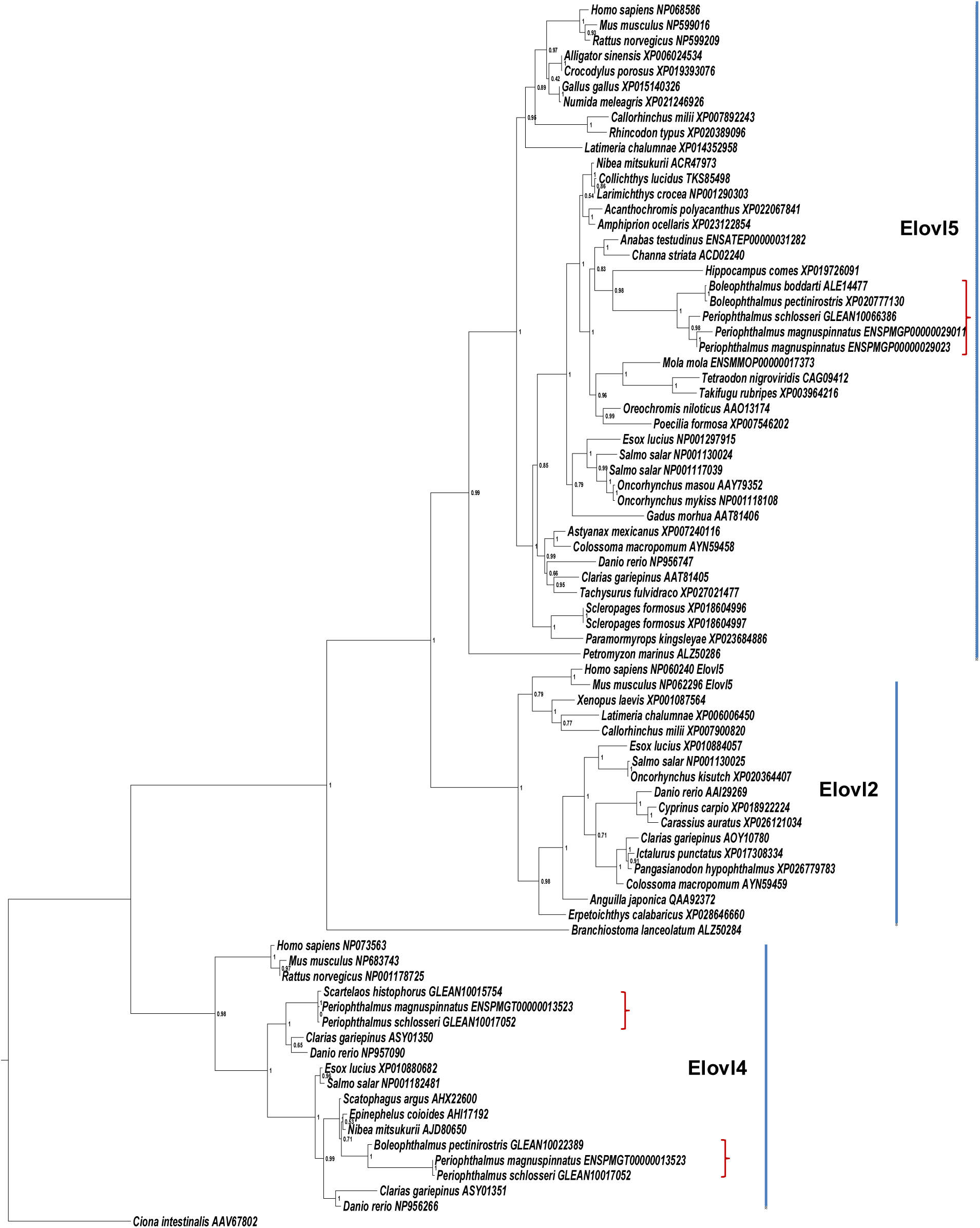
Maximum Likelihood Phylogenetic analysis, using Abayes as support, of 80 Elovl elongase amino acid sequences from various vertebrate species. The tree was visualised using Fig-Tree v1.3.1 (http://tree.bio.ed.ac.uk/software/figtree/). The eleven Elovl sequences from different mudskipper species are indicated. These analysis were conducted using the NGPhylogeny.fr integrative web service.

### Functional characterisation of *B. boddarti Fads2* and *Elovl*

*S. cerevisiae* inserted with PYES2 vector without the ORF insert contained 16:0, 16:1n-7, 18:0 and 18:1n-9, fatty acids which are endogenous in the yeast (Fig 5). Yeast transformed with *B. boddarti Fads2* cultured with addition of LNA and LA showed presence of the C18:4n-3 and C18:3n-6, the respective Δ6 desaturation products of the two substrates, albeit at low activity levels (Table 3). As for the rest of the substrates, no desaturation product was obtainedTherefore, the cloned *B. boddarti* Fads is a unifunctional Δ6 Fads2 with low activities.

**Fig. 5.**
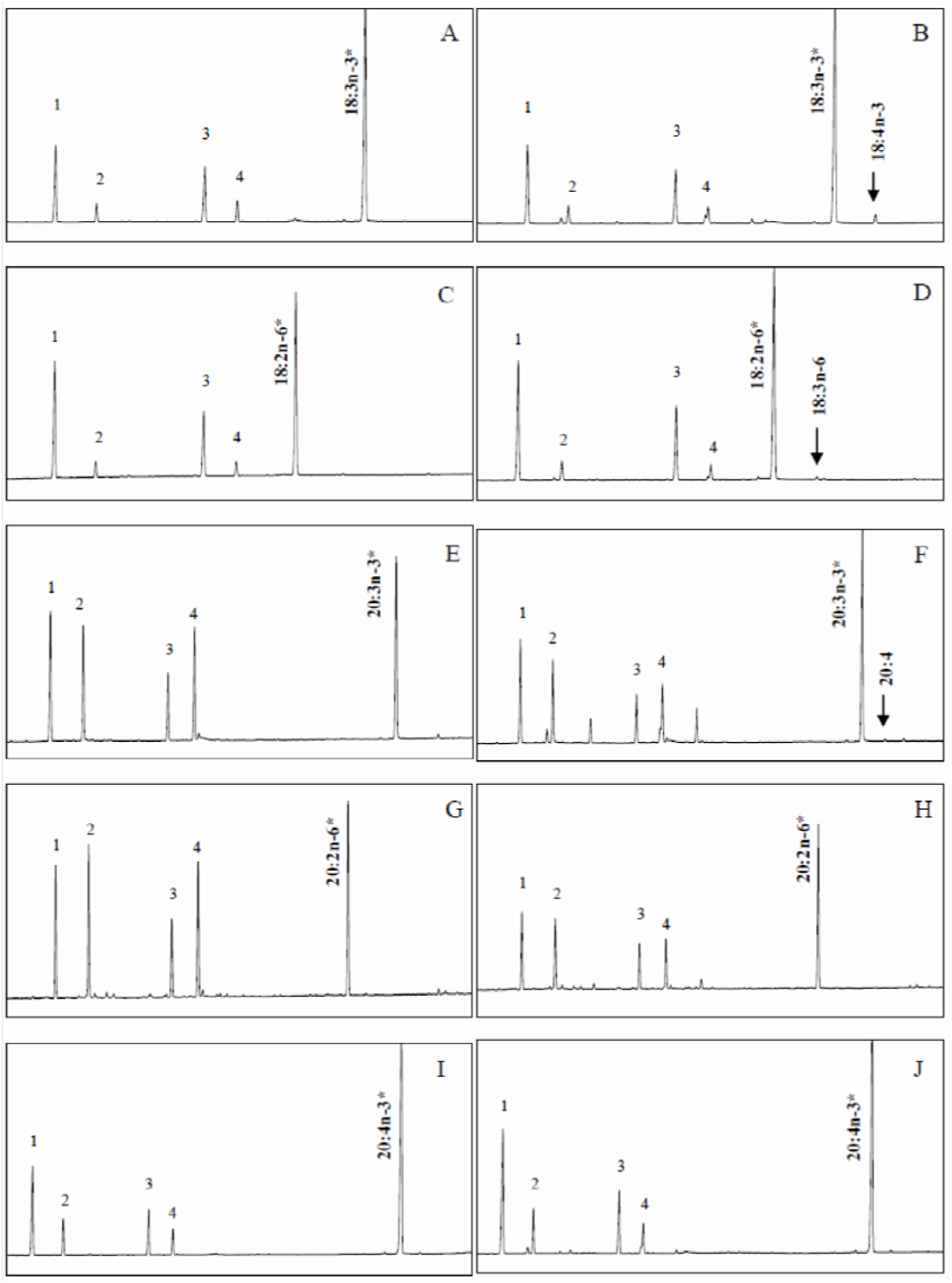
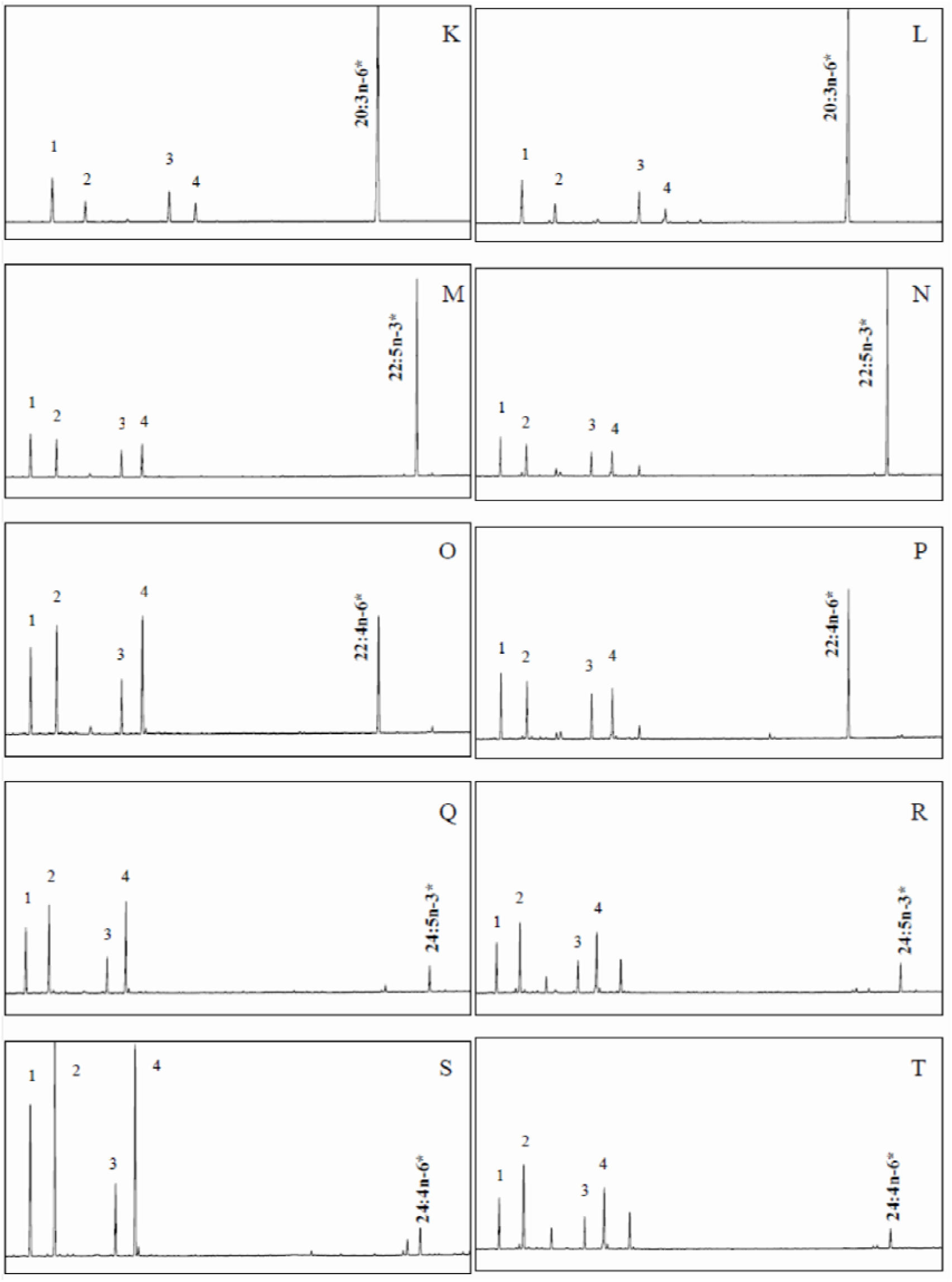
Fatty acid profile of *Saccharomyces cerevisiae* transformed with *Boleophthalmus boddarti fads* (ALE14476) and grown in the presence of substrates for Δ6 desaturation: 18:3n-3 (B), 18:2n-6 (D), 24:5n-3 (R) and 24:4n-6 (T); Δ8 desaturation: 20:3n-3 (F) and 20:2n-6 (H); Δ5 substrates: 20:4n-3 (J) and 20:3n-6 (L) and Δ4 substrates: 22:5n-3 (N) and 22:4n-6 (P). Panels A, C, E, G, I, K, M, O, Q and S are fatty acid profile of *S. cerevisiae* transformed with empty pYES2 vector and incubated with the substrate similar to the corresponding right panel. The first four peaks represent the major endogenous fatty acids of yeast, namely 16:0 (1), 16:1n-7 (2), 18:0 (3) and 18:1n-9 (4). Asterisks indicate exogenously added fatty acid substrates.

**Table 3:** Functional characterization of *Boleophthalmus boddarti fads2* (ALE 14476.1) and *elovl5* (ALE14471.1) via heterologous expression in yeast *Saccharomyces cerevisiae*. The results are expressed as percentage of the fatty acid substrates converted to respective products. Values are presented as Mean ± SEM (*n* = 3).

| Fatty Acid Substrate | Fatty Acid Product | Conversion (%) | Activity |
| --- | --- | --- | --- |
| *B. boddarti fads2* (ALE 14476.1) |  |  |  |
| 18:3n-3 | 18:4n-3 | 3.1 ± 0.1 | $\Delta 6$ |
| 18:2n-6 | 18:3n-6 | 1.1 ± 0.0 | $\Delta 6$ |
| 20:3n-3 | 20:4n-3 | ND | $\Delta 8$ |
| 20:2n-6 | 20:3n-6 | ND | $\Delta 8$ |
| 20:3n-3 | 20:4* | 0.6 ± 0.1 | $\Delta 5$ / $\Delta 6$ |
| 20:2n-6 | 20:3* | ND | $\Delta 5$ / $\Delta 6$ |
| 20:4n-3 | 20:5n-3 | ND | $\Delta 5$ |
| 20:3n-6 | 20:4n-6 | ND | $\Delta 5$ |
| 22:5n-3 | 22:6n-3 | ND | $\Delta 4$ |
| 22:4n-6 | 22:5n-6 | ND | $\Delta 4$ |
| 24:5n-3 | 24:6n-3 | ND | $\Delta 6$ |
| 24:4n-6 | 24:5n-6 | ND | $\Delta 6$ |
| *B. boddarti elovl5* (ALE14471.1) |  |  |  |
| 18:3n-3 | 20:3n-3 | 39.5 ± 2.6 | $C_{18} \rightarrow C_{20}$ |
| | 22:3n-3 | 8.8 ± 0.7 | $C_{20} \rightarrow C_{22}$ |
|  | Total | 48 |  |
| 18:2n-6 | 20:2n-6 | 33.1 ± 1.6 | $C_{18} \rightarrow C_{20}$ |
| | 22:2n-6 | 5.9 ± 0.3 | $C_{20} \rightarrow C_{22}$ |
|  | Total | 39 |  |
| 18:4n-3 | 20:4n-3 | 59.2 ± 5.6 | $C_{18} \rightarrow C_{20}$ |
| | 22:4n-3 | 22.5 ± 4.5 | $C_{20} \rightarrow C_{22}$ |
|  | Total | 82 |  |
| 18:3n-6 | 20:3n-6 | 60.5 ± 2.4 | $C_{18} \rightarrow C_{20}$ |
| | 22:3n-6 | 14.1 ± 0.7 | $C_{20} \rightarrow C_{22}$ |
|  | Total | 75 |  |
| 20:5n-3 | 22:5n-3 | 54.5 ± 1.3 | $C_{20} \rightarrow C_{22}$ |
| | 24:5n-3 | 10.2 ± 0.3 | $C_{22} \rightarrow C_{24}$ |
|  | Total | 75 |  |
| 20:4n-6 | 22:4n-6 | 56.1 ± 2.3 | $C_{20} \rightarrow C_{22}$ |
| | 24:4n-6 | 11.4 ± 0.7 | $C_{22} \rightarrow C_{24}$ |
|  | Total | 67 |  |
| 22:5n-3 | 24:5n-3 | 18.6 ± 2.4 | $C_{22} \rightarrow C_{24}$ |
| 22:4n-6 | 24:4n-6 | 7.0 ± 1.0 | $C_{22} \rightarrow C_{24}$ |

\*Non-methylene interrupted FA (20:4 could be $\Delta^{5,11,14,17}$ 20:4 or $\Delta^{6,11,14,17}$ 20:4 while 20:3 could be $\Delta^{5,11,14,17}$ 20:3 or $\Delta^{6,11,14,17}$ 20:3).

Yeast transformed with *B. boddarti* Elovl5 and incubated with either of the C18 PUFA substrates showed elongation into C20 and C22 PUFA products (Fig 6). C20 PUFA substrates were also elongated to C22 and C24 products (Table 3). Incubation with 22:5n-3 or 22:4n-6 also yielded C24:5n-3 (18.6 ± 2%) and C24:4n-6 (7.0 ± 1%), respectively. Collectively, these results confirm the cloned *B. boddarti* elongase is an Elovl5 elongase with a broad range of substrate elongation capacity. The conversion activities of C18 and C20 PUFA are higher than the C22 substrates. The addition of any of the tested PUFA substrates tested to the culture medium of wild type yeast did not yield any desaturation and elongation products, validating the origin of PUFA products from desaturation or elongation of yeast transformed with *B. boddarti* Fads/Elovl.

**Fig. 6.**
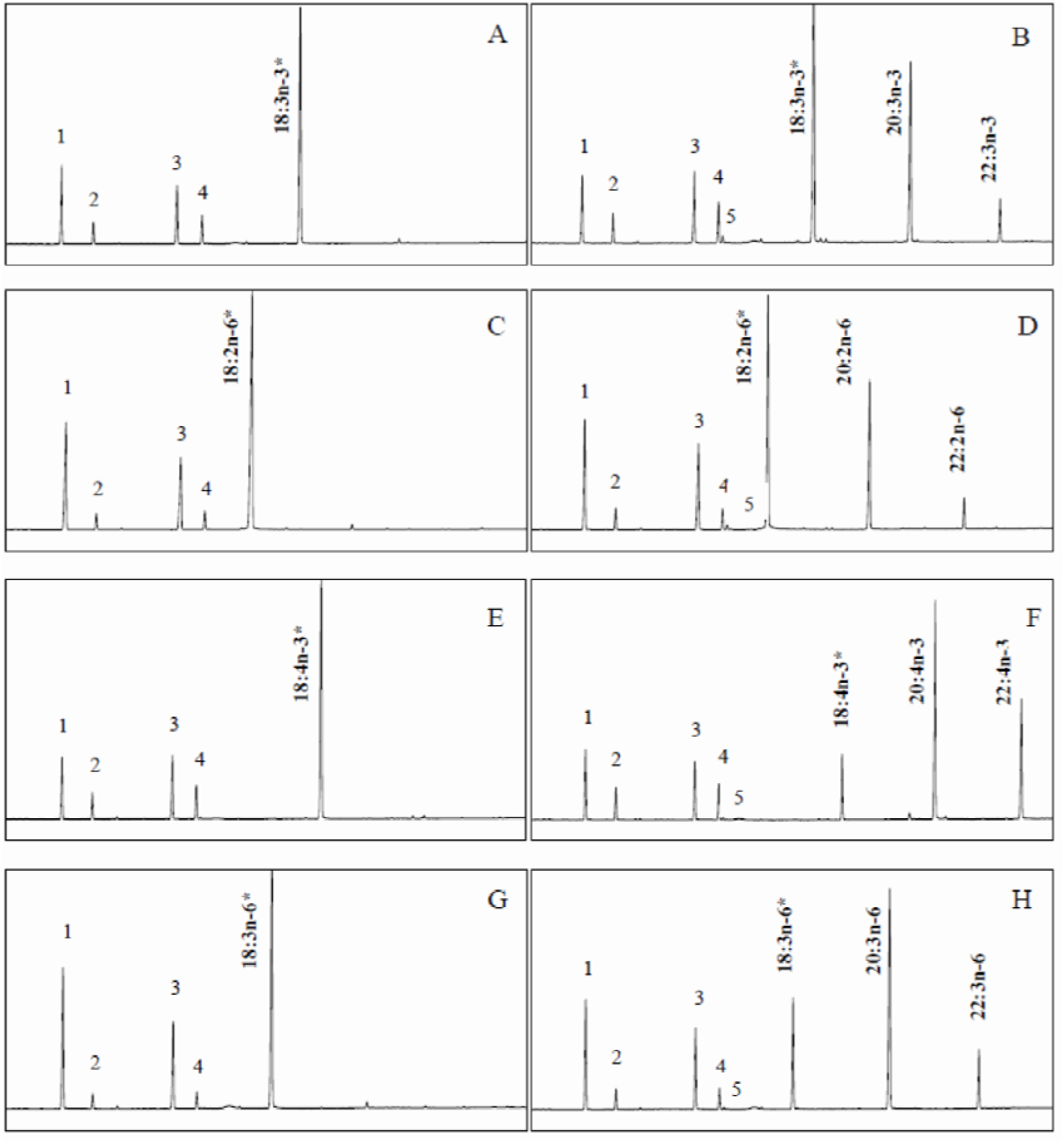
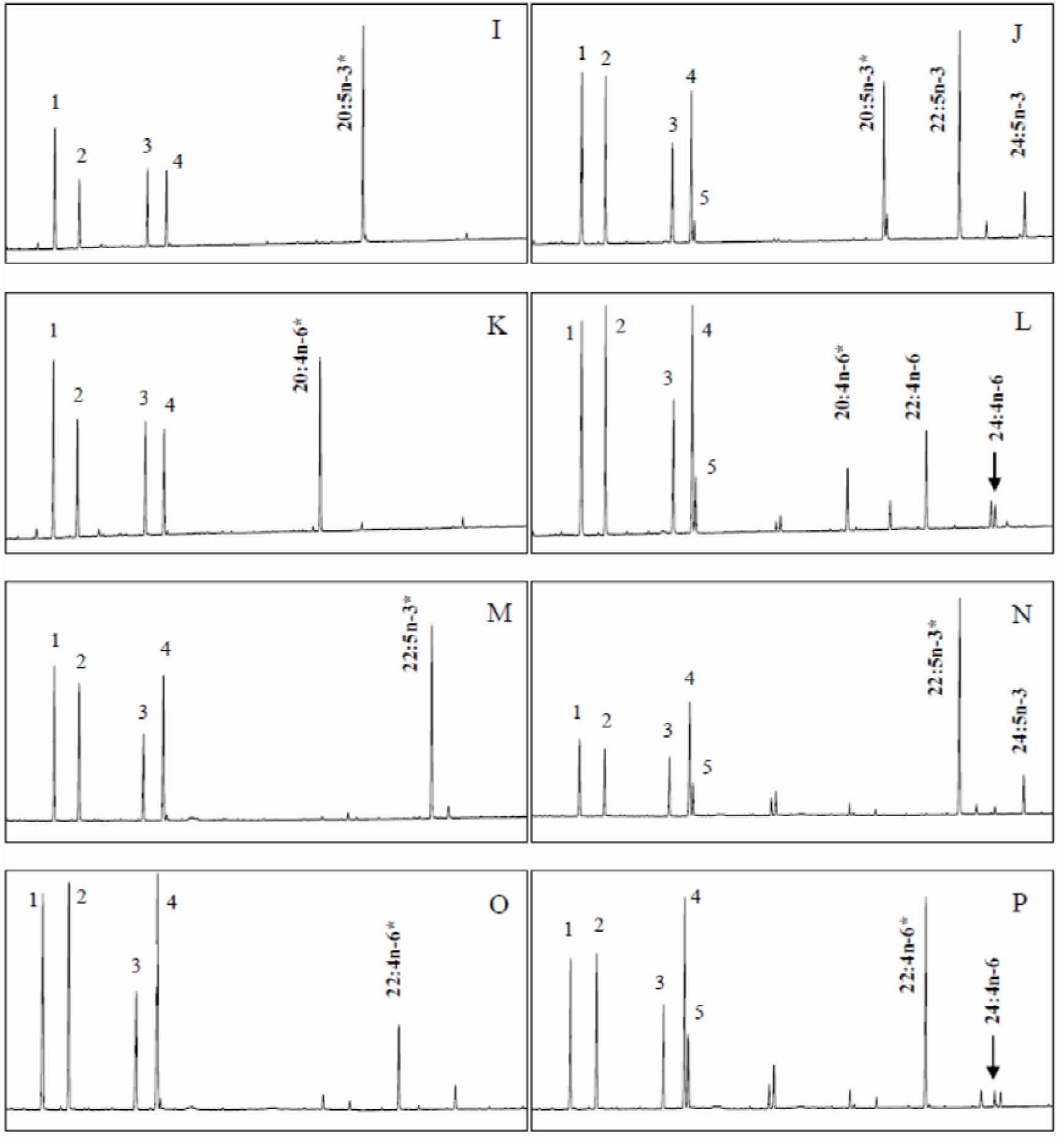
Fatty acid profile of *Saccharomyces cerevisiae* transformed with *Boleophthalmus boddarti elovl* (ALE14477) and grown in the presence of substrates grown in the presence of substrates 18:3n-3 (B), 18:2n-6 (D), 18:4n-3 (F), 18:3n-6 (H), 20:5n-3 (J), 20:4n-6 (L), 22:5n-3 (N) and 22:4n-6 (P). Panels A, C, E, G, I, K, M and O are fatty acid profile sof *S. cerevisiae* transformed with empty pYES2 vector and incubated with the substrate similar to the corresponding right panel. Peaks 1-4 are as described in Fig. 5. Peak 5 corresponds to 18:1n-7 arising from the elongation of the yeast endogenous 16:1n-7. Asterisks indicate exogenously added fatty acid substrates.

### Tissue distribution of *B. boddarti Δ6 Fads2* and *Elovl5*

Among the analysed tissues, brain showed the highest expression of fads2, although statistically, the expression was similar with liver. The remaining tissues showed statistically similar expression levels (Fig. 7). As for *elovl5*, highest level of transcript was observed in intestine.

**Fig 7.**
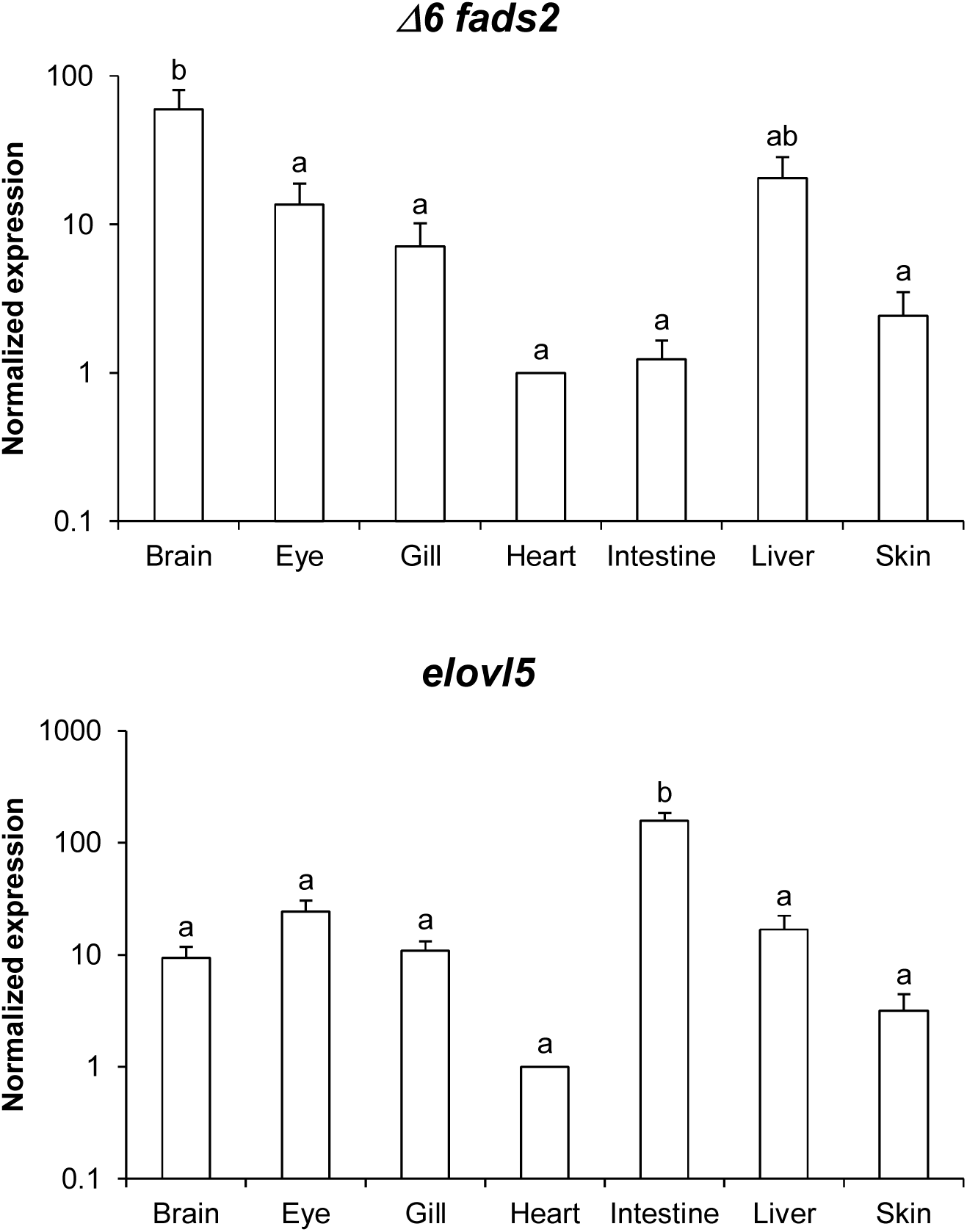
Tissue distribution profile of *fads2* and *elovl5* in adult *Boleophthalmus boddarti* tissues. Expression levels were normalized to a housekeeping gene *elongation factor 1α*. Values shown are mean + SEM (n=3 per tissue). One-way ANOVA followed by Tukey’s post hoc test was performed, with different alphabets indicating a significant difference (P < 0.05).

### Effect of P57Q mutation on elongation activities *of B. boddarti Elovl5*

Findings from the *in vitro* elongation assay revealed the *B. boddarti elovl5* is capable of elongating C22 PUFA substrates, in addition to C18 and C22 substrates typically elongated by teleost Elovl5. We identified a glutamine (Q) residue that seems to be conserved in Elovl5 and appears to be substituted by a proline (P) residue in Elovl2 from various bony fish and the elephant shark *Callorhinchus milii* (Supplementary 1). Interestingly, at this site, the *B. boddarti* Elovl5 possess a P instead of Q. This is also recapitulated in the Elovl5 of *Siganus caniculatus*, which also possess C22 elongation activities (Monroig et al., 2012). Intriguingly, the *elovl2* of the sea lamprey *Petromyzon marinus* possess a Q instead of P and is unable to elongate C22 PUFA substrates (Monroig et al., 2016). Using site-directed mutagenesis, the P residue was replaced with Q in the *B. boddarti* Elovl5. Comparing with the wild type *B. boddarti* Elovl5, the mutated elongase resulted in lower conversion of C22:5n-3, EPA and LNA (Table 4). There was a significant increase of elongation of ARA to C22:4n-6.

**Table 4:** Functional characterization of *Boleophthalmus boddarti elovl5* (ALE14471.1) and *B boddarti* with mutation P-Q at 57 via heterologous expression in yeast *Saccharomyces cerevisiae*. The results are expressed as percentage of the fatty acid substrates converted to elongated products. Values are presented as Mean ± SEM (*n* = 3). * represent significant difference between two clones (student T test, P < 0.05)

| Fatty acid substrate | Fatty acid product | Conversion (%) |  | Activity |
| --- | --- | --- | --- | --- |
|  |  | *B. boddarti elovl5* | *B. boddarti elovl5* PQ |  |
| 18:3n-3 | 20:3n-3 | 39.0 ± 0.2 | 29.1 ± 0.4* | C<sub>18</sub> → C<sub>20</sub> |
| 18:2n-6 | 20:2n-6 | 31.2 ± 0.4 | 31.9 ± 0.4 | C<sub>18</sub> → C<sub>20</sub> |
| 18:4n-3 | 20:4n-3 | 58.8 ± 0.5 | 56.3 ± 0.1 | C<sub>18</sub> → C<sub>20</sub> |
| 18:3n-6 | 20:3n-6 | 62.1 ± 0.7 | 60.1 ± 1.0 | C<sub>18</sub> → C<sub>20</sub> |
| 20:5n-3 | 22:5n-3 | 59.9 ± 0.5 | 51.2 ± 0.4* | C<sub>20</sub> → C<sub>22</sub> |
| 20:4n-6 | 22:4n-6 | 61.0 ± 0.4 | 73.7 ± 0.4* | C<sub>20</sub> → C<sub>22</sub> |
| 22:5n-3 | 24:5n-3 | 16.3 ± 1.1 | 13.6 ± 0.7* | C<sub>22</sub> → C<sub>24</sub> |
| 22:4n-6 | 24:4n-6 | 8.4 ± 1.0 | 7.6 ± 0.1 | C<sub>22</sub> → C<sub>24</sub> |

### Fatty acid composition of mudskipper tissues

DHA concentration was highest in the two neuron-rich organs, eye and brain (Table 5). In addition, there was also a substantial level of DHA deposition in muscle tissue. As for EPA, high concentrations were obtained in gill, gonad, intestine and liver. Highest deposition of ARA was in intestine and muscle. Tissues with highest composition of LNA and LA were gonad and liver, respectively. Saturated fatty acids were abundant in heart, while total monounsaturated fatty acids were highest in brain tissue.

**Table 5:** Fatty acid composition (% of total fatty acids) of various tissues of *Boleophthalmus boddarti*. SFA = Saturated fatty acid; MUFA = Monounsaturated fatty acid; PUFA = Polyunsaturated fatty acid. The results are presented as the means ± SEM (n = 3).

| Fatty acid | Tissues |  |  |  |  |  |  |  |
| --- | --- | --- | --- | --- | --- | --- | --- | --- |
|  | Brain | Eye | Gill | Gonad | Heart | Intestine | Liver | Muscle |
| 10:0 | 0.55 ± 0.02 | 0.75 ± 0.01 | 0.02 ± 0.00 | 0.02 ± 0.01 | 1.46 ± 0.02 | ND | ND | 0.34 ± 0.05 |
| 11:0 | 0.21 ± 0.04 | 0.61 ± 0.01 | 0.03 ± 0.01 | 0.03 ± 0.01 | 1.05 ± 0.01 | ND | 0.02 ± 0.00 | 0.19 ± 0.02 |
| 12:0 | 0.67 ± 0.01 | 0.99 ± 0.01 | 0.04 ± 0.02 | 0.04 ± 0.01 | 2.31 ± 0.06 | ND | 0.03 ± 0.00 | 0.23 ± 0.06 |
| 13:0 | ND | ND | 0.16 ± 0.02 | 0.16 ± 0.00 | ND | ND | 0.21 ± 0.01 | 0.32 ± 0.29 |
| 14:0 | 0.18 ± 0.02 | 0.60 ± 0.02 | 2.69 ± 0.06 | 2.77 ± 0.04 | 0.70 ± 0.02 | 2.17 ± 0.08 | 2.52 ± 0.06 | 0.53 ± 0.01 |
| 15:0 | 0.57 ± 0.01 | 1.95 ± 0.01 | 5.72 ± 0.08 | 5.96 ± 0.08 | 2.07 ± 0.02 | 3.55 ± 0.04 | 9.40 ± 0.17 | 3.30 ± 0.03 |
| 16:0 | 16.35 ± 0.01 | 24.94 ± 0.12 | 25.84 ± 0.13 | 27.33 ± 0.44 | 38.52 ± 0.08 | 24.48 ± 0.61 | 24.84 ± 0.54 | 20.28 ± 0.70 |
| 17:0 | 1.51 ± 0.01 | 1.68 ± 0.04 | 2.30 ± 0.01 | 2.49 ± 0.08 | 1.84 ± 0.00 | 2.57 ± 0.16 | 3.78 ± 0.05 | 2.33 ± 0.07 |
| 18:0 | 16.44 ± 0.00 | 19.82 ± 0.03 | 9.01 ± 0.04 | 8.62 ± 0.64 | 35.40 ± 0.02 | 11.87 ± 0.37 | 6.72 ± 2.08 | 15.47 ± 0.98 |
| 20:0 | 0.24 ± 0.02 | 0.23 ± 0.01 | 0.03 ± 0.01 | 0.03 ± 0.01 | 0.28 ± 0.03 | 0.11 ± 0.01 | 0.33 ± 0.01 | 0.22 ± 0.06 |
| 24:0 | 8.44 ± 0.06 | 2.52 ± 0.05 | 0.15 ± 0.00 | 0.16 ± 0.00 | ND | 4.01 ± 2.31 | 0.23 ± 0.00 | 1.30 ± 0.50 |
| **$\Sigma$ SFA** | **45.15 ± 0.02** | **54.08 ± 0.23** | **45.99 ± 0.12** | **47.59 ± 0.00** | **83.64 ± 0.10** | **48.76 ± 1.05** | **48.09 ± 1.37** | **44.51 ± 1.44** |
| 14:1 | 0.15 ± 0.02 | 0.16 ± 0.01 | 0.01 ± 0.01 | 0.01 ± 0.00 | 0.29 ± 0.04 | ND | 0.08 ± 0.00 | 0.41 ± 0.08 |
| 15:1 | 0.51 ± 0.02 | 1.29 ± 0.02 | 0.18 ± 0.03 | 0.20 ± 0.03 | 1.61 ± 0.05 | ND | 0.28 ± 0.02 | 1.92 ± 0.92 |
| 16:1 | 2.03 ± 0.02 | 2.47 ± 0.01 | 10.79 ± 0.07 | 11.05 ± 0.12 | 1.92 ± 0.07 | 9.30 ± 0.12 | 10.12 ± 0.21 | 2.54 ± 0.06 |
| 17:1 | 6.65 ± 0.00 | 5.41 ± 0.04 | 4.06 ± 0.01 | 1.95 ± 0.16 | 3.30 ± 0.01 | 2.87 ± 0.23 | 3.43 ± 0.06 | 5.34 ± 0.41 |
| 18:1n-9 | 21.06 ± 0.02 | 9.93 ± 0.01 | 3.84 ± 0.01 | 4.00 ± 0.10 | 2.40 ± 0.03 | 3.97 ± 0.10 | 5.76 ± 0.10 | 4.75 ± 0.05 |
| 20:1n-9 | 0.11 ± 0.01 | 0.13 ± 0.01 | 2.53 ± 0.01 | 1.59 ± 0.06 | 0.67 ± 0.01 | 1.46 ± 0.08 | 1.89 ± 0.10 | 0.48 ± 0.02 |
| 22:1n-9 | 0.29 ± 0.02 | 0.23 ± 0.02 | 0.14 ± 0.12 | 0.28 ± 0.01 | ND | ND | 0.27 ± 0.01 | 1.73 ± 0.02 |
| 24:1n-9 | 0.25 ± 0.00 | 0.57 ± 0.01 | 0.27 ± 0.02 | 0.31 ± 0.04 | 0.20 ± 0.01 | ND | 0.63 ± 0.01 | 0.66 ± 0.01 |
| **$\Sigma$ MUFA** | **31.05 ± 0.08** | **20.18 ± 0.03** | **20.98 ± 0.12** | **19.38 ± 0.08** | **9.82 ± 0.10** | **17.60 ± 0.53** | **22.46 ± 0.48** | **17.84 ± 1.44** |
| 18:3n-3 | 0.10 ± 0.02 | 0.28 ± 0.01 | 1.68 ± 0.00 | 1.73 ± 0.07 | 0.34 ± 0.02 | 1.23 ± 0.06 | 1.56 ± 0.07 | 0.42 ± 0.03 |
| 18:4n-3 | 0.07 ± 0.02 | 0.15 ± 0.03 | 0.07 ± 0.04 | 0.12 ± 0.00 | 0.11 ± 0.03 | ND | 0.20 ± 0.01 | 0.16 ± 0.01 |
| 20:3n-3 | 0.09 ± 0.00 | 0.17 ± 0.02 | 0.31 ± 0.00 | 0.33 ± 0.02 | ND | 0.33 ± 0.02 | 0.25 ± 0.00 | 0.39 ± 0.03 |
| 20:4n-3 | 0.09 ± 0.02 | 0.15 ± 0.02 | 0.84 ± 0.00 | 0.88 ± 0.04 | ND | 0.46 ± 0.04 | 0.61 ± 0.02 | 0.34 ± 0.02 |
| 20:5n-3 | 2.21 ± 0.01 | 3.48 ± 0.03 | 12.15 ± 0.12 | 12.11 ± 0.07 | 1.36 ± 0.02 | 11.60 ± 0.39 | 10.87 ± 0.47 | 8.75 ± 0.05 |
| 22:5n-3 | 0.97 ± 0.02 | 2.65 ± 0.04 | 5.58 ± 0.03 | 5.72 ± 0.07 | 0.57 ± 0.02 | 5.00 ± 0.26 | 3.32 ± 0.12 | 4.47 ± 0.01 |
| 22:6n-3 | 14.90 ± 0.07 | 13.55 ± 0.00 | 6.08 ± 0.12 | 5.60 ± 0.40 | 1.31 ± 0.00 | 6.45 ± 0.20 | 2.97 ± 0.07 | 12.98 ± 0.10 |
| **Σ ω-3 PUFA** | **18.43 ± 0.06** | **20.42 ± 0.14** | **26.72 ± 0.02** | **26.49 ± 0.30** | **3.69 ± 0.03** | **25.06 ± 0.45** | **19.77 ± 0.74** | **27.51 ± 0.04** |
| 18:2n-6 | 0.57 ± 0.15 | 0.61 ± 0.05 | 2.53 ± 0.01 | 2.62 ± 0.10 | 0.67 ± 0.01 | 2.29 ± 0.01 | 4.99 ± 0.08 | 1.35 ± 0.02 |
| 18:3n-6 | 0.14 ± 0.01 | 0.14 ± 0.00 | 0.48 ± 0.00 | 0.50 ± 0.02 | ND | 0.41 ± 0.01 | 0.51 ± 0.01 | 0.27 ± 0.01 |
| 20:3n-6 | 1.07 ± 0.01 | 0.39 ± 0.05 | 0.08 ± 0.00 | 0.07 ± 0.03 | ND | 0.13 ± 0.04 | 0.07 ± 0.01 | 0.05 ± 0.01 |
| 20:4n-6 | 2.40 ± 0.01 | 2.99 ± 0.00 | 1.76 ± 0.00 | 1.80 ± 0.05 | 1.49 ± 0.02 | 3.70 ± 0.16 | 1.62 ± 0.04 | 4.95 ± 0.03 |
| **Σ ω-6 PUFA** | **4.18 ± 0.17** | **4.13 ± 0.11** | **4.85 ± 0.00** | **5.03 ± 0.21** | **2.16 ± 0.03** | **6.53 ± 0.11** | **7.19 ± 0.14** | **6.62 ± 0.04** |
| 20:2 | 0.51 ± 0.01 | 1.19 ± 0.01 | 1.45 ± 0.02 | 1.48 ± 0.05 | 0.70 ± 0.00 | 2.05 ± 0.00 | 2.49 ± 0.04 | 3.53 ± 0.08 |
| 22:2 | 0.68 ± 0.02 | ND | 0.01 ± 0.00 | 0.03 ± 0.02 | ND | ND | ND | ND |
| **Σ other PUFAs** | **1.19 ± 0.03** | **1.19 ± 0.01** | **1.46 ± 0.03** | **1.51 ± 0.03** | **0.70 ± 0.01** | **2.05 ± 0.00** | **2.49 ± 0.04** | **3.53 ± 0.08** |
| **n-3/n-6** | **4.41 ± 0.19** | **4.95 ± 0.09** | **5.50 ± 0.01** | **5.27 ± 0.27** | **1.71 ± 0.04** | **3.84 ± 0.01** | **2.75 ± 0.05** | **4.15 ± 0.02** |
| **Σ Total PUFAs** | **23.80 ± 0.10** | **25.73 ± 0.26** | **33.03 ± 0.01** | **33.03 ± 0.05** | **6.55 ± 0.01** | **33.64 ± 0.56** | **29.45 ± 0.89** | **37.66 ± 0.16** |

### In silico discovery of *Fads* and *Elovl* from different mudskipper species

Besides the cloning and characterisation of the *B. boddarti* Fads2 and Elovl5, we also performed an *in silico* search on relevant orthologs from the published genome sequences of *B. pectinirostris*, *S. histophorus, P. schlosseri* and *P. magnuspinnatus*. Search reveals *B. pectinirostris, P. schlosseri* and *P. magnuspinnatus* has a single *fads* and *elovl* orthologs, respectively (Table 2). BLASTn analysis revealed a relatively high similarity (>75%) between these genes and the corresponding Fads/Elovl orthologs from *B. boddarti* and *D. rerio*, respectively. We could not obtain any highly similar hits from *S. histiphorus*. There was also no match (hits below 30% coverage) in any of these species with the zebrafish *elovl2*. As for Elovl4, putative orthologs were mined from all four mudskipper species.

## Discussion

We successfully cloned a full-length *fads* desaturase cDNA from *B. boddarti*, with the predicted amino acid sequence having all the distinctive features of a microsomal fatty acyl front-end desaturase. This include a cytochrome b_5_-like domain at the N-terminal end, which play a role as electron donor during desaturation (Napier et al., 2003). The presence of a heme binding motif HPGG within the domain of the *B. boddarti* Fads sequence is consistent with its crucial role in desaturation (Sayanova et al., 1999). Front end desaturases of many species contain a highly conserved feature of three histidine boxes, HX_3-4_H, HX_2-3_HH, HX_2-3_HH, an active site for desaturation, presumably for formation of diiron complex (Shanklin et al., 1994). *B. boddarti fads* also contain these 3 boxes, with a glutamine residue replacing the first histidine of the third box (QIEHH). This glutamine is conserved in all teleost *fads2* and has been shown to be essential for desaturation activity (Sayanova et al., 2001). In addition, a four amino acid residues FHLQ at 277-280 matching the four residues proposed to confer regioselectivity towards Δ5 and Δ6 desaturation was also present within the third transmembrane region of the *B. boddarti fads* (Lim et al., 2014). Phylogenetic analysis of the *B. boddarti* Fads sequence was done with Fads1 and Fads2 sequences obtained from various taxonomy groups including invertebrates, Chondrichthyes, sarcopterygians and actinopterygians. For the latter group, we obtained sequences from various Teleostei. The cloned *B. boddarti* Fads and the 3 mined Fads sequences from *B. pectinirostris, P. schlosseri* and *P. magnuspinnatus* are clustered together. There was also a clade of well-supported vertebrate Fads1 sequences, which contain orthologs from cartilaginous species (*Callorhinchus milii*), spotted gar (*Lepisosteus oculatus*), Japanese eel (*Anguilla japonica*) and the Senegal bichir (*Polypterus senegalus*). The discovery of functional Fads1 with Δ5 and Fads2 with Δ6 capacities in cartilaginous catshark led to the postulation these enzymes arose from a gene duplication event before the origin of gnathostome (Castro et al., 2012). The Fads1 was retained in Chondrichthyes and early ray-finned fish before its subsequent loss in Osteoglossomorpha and Clupeocephala (Lopes-Marques et al., 2018). The Japanese eel, *Anguilla japonica* is the sole which retained a *fads1* with a Δ5 desaturation activity. As for Fads2, majority of the Teleostei groups not only retain the Δ6 capacity but undergo functionalisation for Δ4, Δ5 and Δ8 desaturation capabilities (Zheng et al., 2004, Zheng et al., 2009a, Tanomman et al., 2013, Fonseca-Madrigal et al., 2014). Many of the characterised Fads2 sequences also have bifunctional desaturation activities (Monroig et al., 2011, Kabeya et al., 2015, Oboh et al., 2017). The Fads from the four mudskipper species shares a common ancestor with Fads2 of various percomorpha fishes.

In addition, a full length Elovl cDNA was also cloned from *B. boddarti*. The sequence possesses all the structural characteristics common for microsomal Elovl family members such as transmembrane domains, conserved motifs, a histidine box and the canonical C-terminal endoplasmic reticulum retention signal (Leonard et al., 2004). In the phylogenetic tree, *B. boddarti* Elovl is placed within a Elovl5 clade, distinct from the Elovl2 and Elovl4 clades. The Elovl5 clade, supported by an elongase from a lamprey species, comprises of several functionally characterised Elovl5 from the relatively ancient salmonids and ostariophysians (Agaba et al., 2004, Agaba et al., 2005, Morais et al., 2009, Carmona-Antonanzas et al., 2013, Ferraz et al., 2019) and a large clade represented by the species-rich modern percomorphs (Kuah et al., 2015, Kabeya et al., 2015, Monroig et al., 2013). This suggests the cloned *B. boddarti* elongase is an ortholog of the Elovl5 family. In addition, orthologs from *B. pectinirostris, P. schlosseri* and *P. magnuspinnatus* were also clustered with the *B. boddarti* Elovl5. While orthologs of Elovl4 were retrieved from these genomes, we did not find any Elovl2 elongases. Both Elovl2 and Elovl5 were hypothesized to arise from genome duplications and neofunctionalization in vertebrate ancestors (Monroig et al., 2016). As opposed to Elovl5, the Elovl2 is absent in most modern teleost, possible due to gene loss event (Morais et al., 2009).

Using a heterologous yeast expression system, we showed that *B. boddarti* Fads2 possess low activity rates towards ALA and LA. Concomitantly, no detectable PUFA product was obtained with the other tested substrates. Taken together, this shows *B. boddarti* possess a unifunctional Fads2 with low Δ6 desaturation capacity. Attempts to characterise Fads2 from several marine carnivorous species including meagre (*Argyrosomus regius*), Atlantic cod (*Gadus morhua*), chu’s croaker (*Nibea coibor*), barramundi (*Lates calcarifer*) and orange-spotted grouper (*Epinephelus coioides*) have reported low (< 10%) rate of desaturation (Tocher et al., 2006, Tu et al., 2012, Monroig et al., 2013, Li et al., 2014, Huang et al., 2017). The low Δ6 desaturation rate of *B. boddarti* Fads2, coupled with the lack of any observed Δ8 and notably, Δ5 desaturation activities suggest an inability for LC-PUFA biosynthesis in this species, akin to those observed in many marine carnivorous species. In addition, there was no Δ6 desaturation of the C24 substrates, which also rules out the capacity to convert EPA to DHA.

Contrary to Fads2, *B. boddarti* Elovl5 demonstrate higher conversion rates, showing the capacity to elongate C18, C20 and C22 substrates. Therefore, *B. boddarti* Elovl5 appear to fulfil all the elongation requirements for production of DHA and ARA from C18 PUFA. In most teleost, the Elovl5 elongates C18 and C20 PUFA at much higher rate than C22 PUFA, with the latter often at conversion rate of below 5%. While the capacity to elongate C22 PUFA seemed exclusive to Elovl2 rather than Elovl5, this paralog has only been isolated and characterised from salmonids and several clupeocephalan species (Agaba et al., 2004, Morais et al., 2009, Oboh et al., 2016, Machado et al., 2018, Ferraz et al., 2019). To date, *elov5* from two euryhaline herbivorous species, spotted scat (*Scatophagus argus*) and rabbitfish (*Siganus canaliculatus*) have been reported to have moderate but measurable elongation capacity of C22 substrates (Monroig et al., 2012, Xie et al., 2016). *B. boddarti* and both these species occupy a lower trophic level than the marine piscivorous finfish, which most likely explain the necessity for a broader range of PUFA elongation. From the perspective of the LC-PUFA biosynthesis pathway, the capacity to elongate C22:5n-3 to C24:5n-3 means a potential use of the ‘Sprecher pathway’ for biosynthesis of DHA from EPA. However, the inability of *B. boddarti* Fads2 to carry out the Δ6 conversion of 24:5n-3 to 24:6n-3 does not complement the C22 elongation ability. This is in contrast with more basal Teleostei clade such as Cypriniformes, Siluriformes and Salmoniformes, where the presence of an Elovl2 is supported by the Δ6 desaturation of C24:5n-3 in Fads2 (Oboh et al., 2016).

A complete elucidation of the LC-PUFA biosynthesis pathway in any given species requires the understanding for the full spectrum of functional capacities of the Fads and Elovl enzymes. Therefore, despite the presence of a single Elovl5 which fulfils all the elongation steps required for the conversion of C18 PUFA to LC-PUFA, the lack of Δ5 and ‘Sprecher pathway’ Δ6 desaturation activities in the *B. boddarti fads2* indicated limited capacity for endogenous LC-PUFA biosynthesis in this species. Although the possibility of *B. boddarti* having other Fads2 with complementing desaturation cannot be ruled out entirely, *in-silico* search on the genomes of *B. pectinirostris*, *P. schlosseri* and *P. magnuspinnatus* yielded only a single *fads2* for each respective species. Functional characterisation of these orthologs will determine if differences in conversion rate, substrate specificity and regioselectivity exists between these species, which also occupy different niches. Works on many marine teleost have also reported deficient LC-PUFA biosynthesis pathways, despite the possession of functional Elovl5 in these species. Atlantic cod, orange grouper, meagre and *B. boddarti* are hampered by having Fads2 with low desaturation activities for all substrates, while species such as cobia and ballan wrasse are limited by the inability to carry out Δ5 desaturation (Agaba et al., 2005, Tocher et al., 2006, Zheng et al., 2009a, Monroig et al., 2013, Li et al., 2014, Kabeya et al., 2018). Elsewhere, two pufferfish species (Tetraodontiformes) were reported to be missing *fads*-like orthologs in their genome (Leaver et al., 2008). It is intriguing why diminished capacity to biosynthesis LC-PUFA in the marine teleost repeatedly involves the compromise of Fads2 desaturation while the elongation capacity of Elovl5 seemed to remain intact. A possible reason for low desaturation activities in Fads2 could be due to inferior expression at the mRNA level. Elsewhere, work on the promoter region of species with low Fads2 activities showed the absence of critical binding site which are crucial for basal expression in species with notable expression levels (Zheng et al., 2009b, Xie et al., 2018).

Collective studies have lend support to the theory that new or additional functions within Fads2 is speculated to be driven by habitat (freshwater vs marine), trophic level and trophic ecology of a particular species (Navarro-Guillén et al., 2014, Kuah et al., 2016). Hence, marine teleost occupying higher trophic levels, in habitats lush with LC-PUFA rich prey could render the biosynthesis pathway inconsequential, leading to the localised loss of Fads activities. Recently, a marine herbivorous teleost with limited dietary LC-PUFA intake was found to have comparable Fads2 functions with carnivorous representatives of closely related species, which led to the suggestion that phylogenetic position could also be responsible for subfunctionalisation of Fads2 in teleost (Garrido et al., 2019). *B. boddarti* is designated mostly as algae and diatom feeder, with the latter containing significant levels of LC-PUFA (Khoo, 1996, Ravi, 2013). This species is not an obligatory herbivore as polychaetes, nematodes, teleost eggs and detritus have been found in stomachs of captured fishes (Khoo, 1996, Ravi, 2013). Among factors leading to dietary shift in *B. boddarti* are monsoon season and ontogenic development (Ravi, 2013). Therefore, the wide-range of available natural diets very likely provide sufficient supply of LC-PUFA to fulfil the requirements of mudskipper, rendering the biosynthesis pathway inconsequential. Existing literature also indicates the biosynthesis and accumulation of LC-PUFA at various trophic levels of the mangrove environment (Hall et al., 2006, Jaseera et al., 2019). A current concern is the prediction of reduced LC-PUFA production in phytoplankton due ocean warming and its downstream impact on higher consumers with limited LC-PUFA biosynthesis capacity, relying mainly on food webs as source (Hixson and Arts, 2016, Vagner et al., 2019).

The ability of the *B. boddarti* Elovl5 to elongate C22 substrates provide the opportunity to determine a particular amino acid residue that could be key for substrate specificity or optimal enzymatic activity. We speculate that the residue P at position 57 of the *B. boddarti* Elov5, which is identical with teleost Elovl2 elongase from multiple species, could be imperative for the elongation of C22 PUFA substrate. While replacement of P with Q did reduce the conversion percentage of C22:5n-3 to C24:5n-3, there was also an unexpected decrease in the elongation of C18 and C20 substrates as well. Therefore, this particular residue is likely to be important for elongation of C18, C20 and C22 substrates. Previously, a cysteine (C) residue was regarded as essential for elongation of C22 PUFA elongation in rat Elovl2 (Gregory et al., 2013). An attempt to substitute a tryptophan (W) residue with C at the equivalent position in sea lamprey Elovl2 incapable of C22 yielded a small but measurable product of C22 PUFA elongation, (Monroig et al., 2016). In relation to this, both Elovl5 elongases of *B. boddarti* and *S. caniculatus* have the ability to elongate C22 substrates despite having a W residue at this position. Collectively, all these findings suggest there are yet to be discovered amino acid residue essential for specificity towards C22 PUFA substrates.

The notable presence of *B. boddarti fads2* transcript in brain tissue is similar to findings from many marine and freshwater species (Monroig et al., 2013, Ren et al., 2013, Tanomman et al., 2013, Wang et al., 2014, Kuah et al., 2016, Janaranjani et al., 2018, Xie et al., 2018). Given the importance of DHA in neuronal-rich tissues such as brain and eye, having the endogenous capacity for LC-PUFA biosynthesis is advantageous when supply from dietary intake is insufficient. As for *B. boddarti elovl5*, although expression can be found in brain, the level in intestine, a known endodermal organ for LC-PUFA biosynthesis activities was also significant. This is comparable to expression patterns in orange-spotted grouper and meagre, where brain was the tissue with highest level of *fads2* while liver or intestines are major sites for *elovl5* (Monroig et al., 2013, Li et al., 2014, Li et al., 2016). The detection of fads2 in B. boddarti gill tissues are similar to findings in Japanese eel and Atlantic salmon (Wang et al., 2014, Lemmetyinen et al., 2013). Salmonid gill filaments produce a wide range of eicosanoid derivatives important for regulation of water permeability in epithelial layers by modulating the ion and electrolyte balance (Knight et al., 1995). In gills of European seabass, desaturation activity was suggested to be an mechanism to adjust gill PUFA composition as response to water temperature changes (Skalli et al., 2006). The *fads2* and *elovl5* in mudskipper gill could be necessary for maintaining the right balance of LC-PUFA composition crucial to support amphibious lifestyle.

Among all tissues, highest levels of DHA were detected in eye and brain which reassert the importance of n-3 LC-PUFA for neuronal activities. An important adaptation for terrestrial conquest in mudskippers is efficient aerial vision (Sayer, 2005). Therefore, besides the reduction of ultraviolet light-related retinal damage, improvement of color vision, and other morphological adaptations, the ability to maintain sufficient supply of DHA could also be an important (You et al., 2014). The high level of DHA and ARA in *B. boddarti* muscle could potentially be a transferable source of LC-PUFA to terrestrial consumers, as mudskippers are often subjected to predation pressure by land predators (Swanson and Gibb, 2004). Significant concentration of EPA and total PUFA was also detected in mudskipper gill tissue (Bystriansky and Ballantyne, 2007). Alterations in teleost gill FA composition to modify membrane permeability in response to fluctuations in salinity and salinity have been reported. In Atlantic salmon, exposure to reducing water temperature resulted in increase of gill total PUFA and EPA content (Liu et al., 2018). Besides EPA, fluctuation in gill ARA content in different water salinity levels was also reported in diadromous teleost (Bystriansky and Ballantyne, 2007, Itokazu et al., 2014). Taken together, our work indicates the importance of LC-PUFA in the gill tissue of mudskipper.

In conclusion, the molecular cloning and functional characterisation of two critical enzymes in the LC-PUFA biosynthesis pathway, coupled with analysis of different tissue fatty acid composition of *B. boddarti* mudskipper were reported in this study. Results show high percentage of EPA and DHA in neuron-rich tissues. Since the capacity for LC-PUFA biosynthesis is impeded by a Fads2 with low and limited desaturation activities, dietary intake is most probably the sole path for this mudskipper species to acquire LC-PUFA to support their physiological requirements.

## Acknowledgements

We thank Universiti Sains Malaysia for funding this research (304/PBIOLOGI/6315180). The postdoctoral fellowship awarded to Dr Kuah Meng Kiat by Universiti Sains Malaysia is also acknowledged.

## References

Agaba, M., Tocher, D.R., Dickson, C.A., Dick, J.R. & Teale, A.J. (2004). Zebrafish cDNA encoding multifunctional fatty acid elongase involved in production of eicosapentaenoic (20:5n-3) and docosahexaenoic (22:6n-3) acids. Mar Biotechnol, 6: 251–261. DOI 10.1007/s10126-003-0029-1

Agaba, M.K., Tocher, D.R., Zheng, X., Dickson, C.A., Dick, J.R. & Teale, A.J. (2005). Cloning and functional characterisation of polyunsaturated fatty acid elongases of marine and freshwater teleost fish. Comp Biochem Phys B, 142: 342–352.

Anisimova, M., Gil, M., Dufayard, J.-F., Dessimoz, C. & Gascuel, O. (2011). Survey of Branch Support Methods Demonstrates Accuracy, Power, and Robustness of Fast Likelihood-based Approximation Schemes. Systematic Biology, 60: 685–699. DOI 10.1093/sysbio/syr041

Arts, M.T., Ackman, R.G. & Holub, B.J. (2001). “Essential fatty acids” in aquatic ecosystems: a crucial link between diet and human health and evolution. Can J Fish Aquat Sci, 58: 122–137. DOI 10.1139/f00-224

Bystriansky, J.S. & Ballantyne, J.S. (2007). Gill Na+-K+-ATPase activity correlates with basolateral membrane lipid composition in seawater-but not freshwater-acclimated Arctic char (*Salvelinus alpinus*). American Journal of Physiology-Regulatory, Integrative and Comparative Physiology, 292: R1043–R1051. DOI 10.1152/ajpregu.00189.2005

Carmona-Antonanzas, G., Tocher, D.R., Taggart, J.B. & Leaver, M.J. (2013). An evolutionary perspective on Elovl5 fatty acid elongase: comparison of Northern pike and duplicated paralogs from Atlantic salmon. BMC Evol Biol, 13: 1–13.

Castro, L.F.C., Monroig, Ó., Leaver, M.J., Wilson, J., Cunha, I. & Tocher, D.R. (2012). Functional desaturase *Fads1 Α5* and *Fads2 Α6* orthologues evolved before the origin of jawed vertebrates. PLoS ONE, 7: e31950. DOI 10.1371/journal.pone.0031950

Castro, L.F.C., Tocher, D.R. & Monroig, O. (2016). Long-chain polyunsaturated fatty acid biosynthesis in chordates: Insights into the evolution of Fads and Elovl gene repertoire. Prog Lipid Res, 62: 25–40. DOI 10.1016/j.plipres.2016.01.001

Clayton, D.A. & Wright, J.M. (1989). Mud-walled territories and feeding behaviour of *Boleophthalmus boddarti* (Pisces: Gobiidae) on the Mudflats of Kuwait. Journal of Ethology, 7: 91–95. DOI 10.1007/bf02350029

Coelho, H., Lopes Da Silva, T., Reis, A., Queiroga, H., Serôdio, J. & Calado, R. (2011). Fatty acid profiles indicate the habitat of mud snails *Hydrobia ulvae* within the same estuary: Mudflats vs. seagrass meadows. Estuarine, Coastal and Shelf Science, 92: 181–187. DOI https://doi.org/10.1016/j.ecss.2011.01.005

De Deckere, E.a.M. (2001). Health Aspects of Fish and n-3 Polyunsaturated Fatty Acids from Plant and Marine Origin. IN Wilson, T. & Temple, N. (Eds.) Nutritional Health. Humana Press.

De Troch, M., Boeckx, P., Cnudde, C., Van Gansbeke, D., Vanreusel, A., Vincx, M. & Caramujo, M.J. (2012). Bioconversion of fatty acids at the basis of marine food webs: insights from a compound-specific stable isotope analysis. Mar Ecol Prog Ser, 465: 53–67.

Ferraz, R.B., Kabeya, N., Lopes-Marques, M., Machado, A.M., Ribeiro, R.A., Salaro, A.L., Ozório, R., Castro, L.F.C. & Monroig, Ó. (2019). A complete enzymatic capacity for long-chain polyunsaturated fatty acid biosynthesis is present in the Amazonian teleost tambaqui, Colossoma macropomum. Comp Biochem Physiol B, 227: 90–97. DOI https://doi.org/10.1016/j.cbpb.2018.09.003

Fonseca-Madrigal, J., Navarro, J.C., Hontoria, F., Tocher, D.R., Martínez-Palacios, C.A. & Monroig, Ó. (2014). Diversification of substrate specificities in teleostei Fads2: characterization of Δ4 and Δ6Δ5 desaturases of *Chirostoma estor*. J Lipid Res, 55: 1408–1419. DOI 10.1194/jlr.M049791

Garrido, D., Kabeya, N., Betancor, M.B., Pérez, J.A., Acosta, N.G., Tocher, D.R., Rodríguez, & Monroig, Ó. (2019). Functional diversification of teleost Fads2 fatty acyl desaturases occurs independently of the trophic level. Sci Rep, 9: 11199. DOI 10.1038/s41598-019-47709-0

Gladyshev, M.I., Arts, M.T. & Sushchik, N.N. (2009). Preliminary estimates of the export of omega-3 highly unsaturated fatty acids (EPA+DHA) from aquatic to terrestrial ecosystems. IN Kainz, M., Brett, M.T. & Arts, M.T. (Eds.) Lipids in Aquatic Ecosystems. New York, NY: Springer New York.

Gregory, M.K., Cleland, L.G. & James, M.J. (2013). Molecular basis for differential elongation of omega-3 docosapentaenoic acid by the rat Elovl5 and Elovl2. J Lipid Res, 54: 2851–2857. DOI 10.1194/jlr.M041368

Guindon, S., Dufayard, J.-F., Lefort, V., Anisimova, M., Hordijk, W. & Gascuel, O. (2010). New Algorithms and Methods to Estimate Maximum-Likelihood Phylogenies: Assessing the Performance of PhyML 3.0. Systematic Biology, 59: 307–321. DOI 10.1093/sysbio/syq010

Guo, F., Bunn, S.E., Brett, M.T. & Kainz, M.J. (2017). Polyunsaturated fatty acids in stream food webs – high dissimilarity among producers and consumers. Freshwater Biol, 62: 1325–1334. DOI doi:10.1111/fwb.12956

Hall, D., Lee, S.Y. & Meziane, T. (2006). Fatty acids as trophic tracers in an experimental estuarine food chain: Tracer transfer. J Exp Mar Biol Ecol, 336: 42–53. DOI https://doi.org/10.1016/j.jembe.2006.04.004

Hastings, N., Agaba, M., Tocher, D.R., Leaver, M.J., Dick, J.R., Sargent, J.R. & Teale, A.J. (2001). A vertebrate fatty acid desaturase with Δ5 and Δ6 activities. Proc Natl Acad Sci USA, 98: 14304–14309. DOI 10.1073/pnas.251516598

Hastings, N., Agaba, M.K., Tocher, D.R., Zheng, X., Dickson, C.A., Dick, J.R. & Teale, A.J. (2004). Molecular cloning and functional characterization of fatty acyl desaturase and elongase cDNAs involved in the production of eicosapentaenoic and docosahexaenoic acids from α-linolenic acid in Atlantic Salmon (*Salmo salar*). Mar Biotechnol, 6: 463–474.

Hixson, S.M. & Arts, M.T. (2016). Climate warming is predicted to reduce omega-3, long-chain, polyunsaturated fatty acid production in phytoplankton. Global Change Biology, 22: 2744–2755. DOI 10.1111/gcb.13295

Huang, Y., Lin, Z., Rong, H., Hao, M., Zou, W., Li, S. & Wen, X. (2017). Cloning, tissue distribution, functional characterization and nutritional regulation of Δ6 fatty acyl desaturase in chu’s croaker *Nibea coibor*. Aquaculture, 479: 208–216. DOI https://doi.org/10.1016/j.aquaculture.2017.05.041

Itokazu, Y., Käkelä, R., Piironen, J., Guan, X.L., Kiiskinen, P. & Vornanen, M. (2014). Gill tissue lipids of salmon (*Salmo salar* L.) presmolts and smolts from anadromous and landlocked populations. Comp Biochem Physiol A, 172: 39–45. DOI https://doi.org/10.1016/j.cbpa.2014.01.020

Janaranjani, M., Mah, M.-Q., Kuah, M.-K., Fadhilah, N., Hing, S.-R., Han, W.-Y. & Shu-Chien, A.C. (2018). Capacity for eicosapentaenoic acid and arachidonic acid biosynthesis in silver barb (*Barbonymus gonionotus):* Functional characterisation of a Δ6/Δ8/Δ5 Fads2 desaturase and Elovl5 elongase. Aquaculture, 497: 469–486. DOI https://doi.org/10.1016/j.aquaculture.2018.08.019

Jaseera, K.V., Kaladharan, P., Vijayan, K.K., Sandhya, S.V., Antony, M.L. & Pradeep, M.A. (2019). Isolation and phylogenetic identification of heterotrophic thraustochytrids from mangrove habitats along the southwest coast of India and prospecting their PUFA accumulation. Journal of Applied Phycology, 31: 1057–1068. DOI 10.1007/s10811-018-1627-7

Jump, D.B. (2002). The biochemistry of n-3 polyunsaturated fatty acids. J Biol Chem, 277: 8755–8758. DOI 10.1074/ibc.R100062200

Kabeya, N., Yamamoto, Y., Cummins, S.F., Elizur, A., Yazawa, R., Takeuchi, Y., Haga, Y., Satoh, S. & Yoshizaki, G. (2015). Polyunsaturated fatty acid metabolism in a marine teleost, Nibe croaker Nibea mitsukurii: Functional characterization of Fads2 desaturase and Elovl5 and Elovl4 elongases. Comp Biochem Phys B, 188: 37–45. DOI 10.1016/j.cbpb.2015.06.005

Kabeya, N., Yevzelman, S., Oboh, A., Tocher, D.R. & Monroig, O. (2018). Essential fatty acid metabolism and requirements of the cleaner fish, ballan wrasse *Labrus bergylta:* Defining pathways of long-chain polyunsaturated fatty acid biosynthesis. Aquaculture, 488: 199–206. DOI https://doi.org/10.1016/j.aquaculture.2018.01.039

Katoh, K., Kuma, K.-I., Toh, H. & Miyata, T. (2005). MAFFT version 5: improvement in accuracy of multiple sequence alignment. Nucleic Acids Research, 33: 511–518. DOI 10.1093/nar/gki198

Katoh, K. & Standley, D.M. (2013). MAFFT Multiple Sequence Alignment Software Version 7: Improvements in Performance and Usability. Mol Biol Evol, 30: 772–780. DOI 10.1093/molbev/mst010

Khoo, K. (1996). Studies of the periophthalmid fishes in Singapore. University of Singapore.

Knight, J., Holland, J.W., Bowden, L.A., Halliday, K. & Rowley, A.F. (1995). Eicosanoid generating capacities of different tissues from the rainbow trout, *Oncorhynchus mykiss*. Lipids, 30: 451–458. DOI 10.1007/bf02536304

Kuah, M.-K., Jaya-Ram, A. & Shu-Chien, A.C. (2015). The capacity for long-chain polyunsaturated fatty acid synthesis in a carnivorous vertebrate: Functional characterisation and nutritional regulation of a Fads2 fatty acyl desaturase with Δ4 activity and an Elovl5 elongase in striped snakehead (*Channa striata*). BBA-Mol Cell Biol Lipids, 1851: 248–260. DOI http://dx.doi.org/10.1016/j.bbalip.2014.12.012

Kuah, M.-K., Jaya-Ram, A. & Shu-Chien, A.C. (2016). A fatty acyl desaturase (fads2) with dual Δ6 and Δ5 activities from the freshwater carnivorous striped snakehead *Channa striata*. Comp Biochem Phys A, 201: 146–155. DOI http://dx.doi.org/10.1016/j.cbpa.2016.07.007

Leaver, M.J., Bautista, J.M., Björnsson, B.T., Jönsson, E., Krey, G., Tocher, D.R. & Torstensen, B.E. (2008). Towards Fish Lipid Nutrigenomics: Current State and Prospects for Fin-Fish Aquaculture. Rev Fish Sci, 16: 73–94. DOI 10.1080/10641260802325278

Lefort, V., Longueville, J.-E. & Gascuel, O. (2017). SMS: Smart Model Selection in PhyML. Mol Biol Evol, 34: 2422–2424. DOI 10.1093/molbev/msx149

Lemmetyinen, J., Piironen, J., Kiiskinen, P.I., Hassinen, M. & Vornanen, M. (2013). Comparison of gene expression in the gill of salmon (*Salmo salar*) smolts from anadromous and landlocked populations. Annales Zoologici Fennici, 50: 16–35.

Lemoine, F., Correia, D., Lefort, V., Doppelt-Azeroual, O., Mareuil, F., Cohen-Boulakia, S. & Gascuel, O. (2019). NGPhylogeny.fr: new generation phylogenetic services for non-specialists. Nucleic Acids Research, 47: W260–W265. DOI 10.1093/nar/gkz303

Leonard, A.E., Pereira, S.L., Sprecher, H. & Huang, Y.-S. (2004). Elongation of long-chain fatty acids. Prog Lipid Res, 43: 36–54. DOI http://dx.doi.org/10.1016/S0163-7827(03)00040-7

Li, S., Mai, K., Xu, W., Yuan, Y., Zhang, Y. & Ai, Q. (2014). Characterization, mRNA expression and regulation of Δ6 fatty acyl desaturase (FADS2) by dietary n - 3 long chain polyunsaturated fatty acid (LC-PUFA) levels in grouper larvae (*Epinephelus coioides*). Aquaculture, 434: 212–219. DOI http://dx.doi.org/10.1016/j.aquaculture.2014.08.009

Li, S., Yuan, Y., Wang, T., Xu, W., Li, M., Mai, K. & Ai, Q. (2016). Molecular Cloning, Functional Characterization and Nutritional Regulation of the Putative Elongase Elovl5 in the Orange-Spotted Grouper (*Epinephelus coioides*). PLOS ONE, 11: e0150544. DOI 10.1371/journal.pone.0150544

Li, Y., Monroig, O., Zhang, L., Wang, S., Zheng, X., Dick, J.R., You, C. & Tocher, D.R. (2010). Vertebrate fatty acyl desaturase with Δ 4 activity. Proc Natl Acad Sci USA, 107: 16840–16845. DOI 10.1073/pnas.1008429107

Lim, Z., Senger, T. & Vrinten, P. (2014). Four amino acid residues influence the substrate chain-length and regioselectivity of *Siganus canaliculatus* Δ4 and Δ5/6 desaturases. Lipids, 49: 357–367. DOI 10.1007/s11745-014-3880-0

Liu, C., Zhou, Y., Dong, K., Sun, D., Gao, Q. & Dong, S. (2018). Differences in fatty acid composition of gill and liver phospholipids between Steelhead trout (*Oncorhynchus mykiss*) and Atlantic salmon (*Salmo salar*) under declining temperatures. Aquaculture, 495: 815–822. DOI https://doi.org/10.1016/j.aquaculture.2018.06.045

Lopes-Marques, M., Kabeya, N., Qian, Y., Ruivo, R., Santos, M.M., Venkatesh, B., Tocher, D.R., Castro, L.F.C. & Monroig, Ó. (2018). Retention of fatty acyl desaturase 1 (fads1) in Elopomorpha and Cyclostomata provides novel insights into the evolution of long-chain polyunsaturated fatty acid biosynthesis in vertebrates. BMC Evol Biol, 18: 157. DOI 10.1186/s12862-018-1271-5

Machado, A.M., Tørresen, O.K., Kabeya, N., Couto, A., Petersen, B., Felício, M., Campos, P.F., Fonseca, E., Bandarra, N., Lopes-Marques, M., Ferraz, R., Ruivo, R., Fonseca, M.M., Jentoft, S., Monroig, Ó., Da Fonseca, R.R. & C. Castro, L.F. (2018). “Out of the Can”: A Draft Genome Assembly, Liver Transcriptome, and Nutrigenomics of the European Sardine, Sardina pilchardus. Genes, 9: 485.

Monroig, O., Li, Y. & Tocher, D.R. (2011). Delta-8 desaturation activity varies among fatty acyl desaturases of teleost fish: High activity in delta-6 desaturases of marine species. Comp Biochem Phys B, 159: 206–213. DOI 10.1016/j.cbpb.2011.04.007

Monroig, Ó., Lopes-Marques, M., Navarro, J.C., Hontoria, F., Ruivo, R., Santos, M.M., Venkatesh, B., Tocher, D.R. & C. Castro, L.F. (2016). Evolutionary functional elaboration of the Elovl2/5 gene family in chordates. Sci Rep, 6: 20510. DOI 10.1038/srep20510

Monroig, O., Rotllant, J., Sanchez, E., Cerda-Reverter, J.M. & Tocher, D.R. (2009). Expression of long-chain polyunsaturated fatty acid (LC-PUFA) biosynthesis genes during zebrafish *Danio rerio* early embryogenesis. BBA-Mol Cell Biol Lipids, 1791: 1093–1101.

Monroig, Ó., Tocher, D.R., Hontoria, F. & Navarro, J.C. (2013). Functional characterisation of a Fads2 fatty acyl desaturase with Δ6/Δ8 activity and an Elovl5 with C16, C18 and C20 elongase activity in the anadromous teleost meagre (*Argyrosomus regius*). Aquaculture, 412-413: 14–22.

Monroig, Ó., Wang, S., Zhang, L., You, C., Tocher, D.R. & Li, Y. (2012). Elongation of long-chain fatty acids in rabbitfish *Siganus canaliculatus:* Cloning, functional characterisation and tissue distribution of Elovl5- and Elovl4-like elongases. Aquaculture, 350-353: 63–70. DOI http://dx.doi.org/10.1016/j.aquaculture.2012.04.017

Morais, S., Castanheira, F., Martinez-Rubio, L., Conceição, L.E.C. & Tocher, D.R. (2012). Long chain polyunsaturated fatty acid synthesis in a marine vertebrate: Ontogenetic and nutritional regulation of a fatty acyl desaturase with Δ4 activity. BBA-Mol Cell Biol Lipids, 1821: 660–671. DOI http://dx.doi.org/10.1016/j.bbalip.2011.12.011

Morais, S., Monroig, O., Zheng, X., Leaver, M. & Tocher, D. (2009). Highly unsaturated fatty acid synthesis in Atlantic Salmon: characterization of ELOVL5- and ELOVL2-like elongases. Mar Biotechnol, 11: 627–639.

Murphy, E.O. & Jaafar, Z. (2017). Taxonomy and systematics review. IN Jaafar, Z. & Murphy, E.O. (Eds.) Fishes out of water: biology and ecology of mudskippers. Boca Raton: Taylor & Francis.

Napier, J.A., Michaelson, L.V. & Sayanova, O. (2003). The role of cytochrome b5 fusion desaturases in the synthesis of polyunsaturated fatty acids. Prostaglandins, Leukotrienes and Essential Fatty Acids, 68: 135–143. DOI https://doi.org/10.1016/S0952-3278(02)00263-6

Navarro-Guillén, C., Engrola, S., Castanheira, F., Bandarra, N., Hachero-Cruzado, I., Tocher, D.R., Conceição, L.E.C. & Morais, S. (2014). Effect of varying dietary levels of LC-PUFA and vegetable oil sources on performance and fatty acids of Senegalese sole post larvae: Puzzling results suggest complete biosynthesis pathway from C18 PUFA to DHA. Comp Biochem Phys B, 167: 51–58. DOI http://dx.doi.org/10.1016/j.cbpb.2013.10.001

Oboh, A., Betancor, M.B., Tocher, D.R. & Monroig, O. (2016). Biosynthesis of long-chain polyunsaturated fatty acids in the African catfish *Clarias gariepinus:* Molecular cloning and functional characterisation of fatty acyl desaturase (fads2) and elongase (elovl2) cDNAs7. Aquaculture, 462: 70–79. DOI http://dx.doi.org/10.1016/j.aquaculture.2016.05.018

Oboh, A., Kabeya, N., Carmona-Antonañzas, G., Castro, L.F.C., Dick, J.R., Tocher, D.R. & Monroig, O. (2017). Two alternative pathways for docosahexaenoic acid (DHA, 22:6n-3) biosynthesis are widespread among teleost fish. Sci Rep, 7: 3889. DOI 10.1038/s41598-017-04288-2

Parenti, L.R. & Jaafar, Z. (2017). The natural distribution of mudskippers. IN Jaafar, Z. & Murphy, E.O. (Eds.) Fishes out of water: biology and ecology of mudskippers. Boca Raton: Taylor & Francis.

Prosper, L.M., Tarik, M., Zainudin, B. & Makoto, T. (2003). Fatty acids in decomposing mangrove leaves: microbial activity, decay and nutritional quality. Mar Ecol Prog Ser, 265: 97–105.

Ravi, V. (2013). Food and feeding habits of the mudskipper, *Boleophthalmus boddarti* (Pallas, 1770) from Pichavaram mangroves, Southeast Coast of India. International Journal of Marine Science, 3: 98–104.

Ren, H.T., Zhang, G.Q., Li, J.L., Tang, Y.K., Li, H.X., Yu, J.H. & Xu, P. (2013). Two Δ6-desaturase-like genes in common carp (*Cyprinus carpio* var. Jian): structure characterization, mRNA expression, temperature and nutritional regulation. Gene, 525: 11–7. DOI 10.1016/j.gene.2013.04.073

Sayanova, O., Beaudoin, F., Libisch, B., Castel, A., Shewry, P.R. & Napier, J.A. (2001). Mutagenesis and heterologous expression in yeast of a plant A6Dfatty acid desaturase. Journal of Experimental Botany, 52: 1581–1585. DOI 10.1093/jexbot/52.360.1581

Sayanova, O., Shewry, P.R. & Napier, J.A. (1999). Histidine-41 of the cytochrome b5 domain of the borage delta6 fatty acid desaturase is essential for enzyme activity. Plant Physiol, 121: 641–6.

Sayer, M.D.J. (2005). Adaptations of amphibious fish for surviving life out of water. Fish and Fisheries, 6: 186–211. DOI 10.1111/j.1467-2979.2005.00193.x

Shanklin, J., Whittle, E. & Fox, B.G. (1994). Eight histidine residues are catalytically essential in a membrane-associated iron enzyme, stearoyl-CoA desaturase, and are conserved in alkane hydroxylase and xylene monooxygenase. Biochemistry, 33: 12787–94.

Skalli, A., Robin, J.H., Le Bayon, N., Le Delliou, H. & Person-Le Ruyet, J. (2006). Impact of essential fatty acid deficiency and temperature on tissues’ fatty acid composition of European sea bass (*Dicentrarchus labrax*). Aquaculture, 255: 223–232. DOI https://doi.org/10.1016/j.aquaculture.2005.12.006

Sprecher, H., Luthria, D.L., Mohammed, B.S. & Baykousheva, S.P. (1995). Reevaluation of the pathways for the biosynthesis of polyunsaturated fatty acids. J Lipid Res, 36: 2471–2477.

Swanson, B.O. & Gibb, A.C. (2004). Kinematics of aquatic and terrestrial escape responses in mudskippers. Journal of Experimental Biology, 207: 4037–4044. DOI 10.1242/jeb.01237

Tanomman, S., Ketudat-Cairns, M., Jangprai, A. & Boonanuntanasarn, S. (2013). Characterization of fatty acid delta-6 desaturase gene in Nile tilapia and heterogenous expression in *Saccharomyces cerevisiae*. Comp Biochem Phys B, 166: 148–156. DOI http://dx.doi.org/10.1016/j.cbpb.2013.07.011

Tocher, D.R., Zheng, X., Schlechtriem, C., Hastings, N., Dick, J. & Teale, A. (2006). Highly unsaturated fatty acid synthesis in marine fish: Cloning, functional characterization, and nutritional regulation of fatty acyl Δ6 desaturase of Atlantic cod (*Gadus morhua* L.). Lipids, 41: 1003–1016.

Tu, W.-C., Cook-Johnson, R., James, M., Mühlhäusler, B., Stone, D.J. & Gibson, R. (2012). Barramundi (*Lates calcarifer*) desaturase with Δ6/Δ8 dual activities. Biotechnol Lett, 34: 1283–1296. DOI 10.1007/s10529-012-0891-x

Vagner, M., Pante, E., Viricel, A., Lacoue-Labarthe, T., Zambonino-Infante, J.-L., Quazuguel, P., Dubillot, E., Huet, V., Le Delliou, H., Lefrançois, C. & Imbert-Auvray, N. (2019). Ocean warming combined with lower omega-3 nutritional availability impairs the cardio-respiratory function of a marine fish. The Journal of Experimental Biology, 222: jeb187179. DOI 10.1242/jeb.187179

Wang, S., Monroig, Ó., Tang, G., Zhang, L., You, C., Tocher, D.R. & Li, Y. (2014). Investigating long-chain polyunsaturated fatty acid biosynthesis in teleost fish: Functional characterization of fatty acyl desaturase (Fads2) and Elovl5 elongase in the catadromous species, Japanese eel *Anguilla japonica*. Aquaculture, 434: 57–65. DOI http://dx.doi.org/10.1016/j.aquaculture.2014.07.016

Xie, D., Chen, F., Lin, S., You, C., Wang, S., Zhang, Q., Monroig, Ó., Tocher, D.R. & Li, Y. (2016). Long-chain polyunsaturated fatty acid biosynthesis in the euryhaline herbivorous teleost *Scatophagus argus:* Functional characterization, tissue expression and nutritional regulation of two fatty acyl elongases. Comp Biochem Phys B, 198: 37–45. DOI http://dx.doi.org/10.1016/j.cbpb.2016.03.009

Xie, D., Fu, Z., Wang, S., You, C., Monroig, O., Tocher, D.R. & Li, Y. (2018). Characteristics of the *fads2* gene promoter in marine teleost *Epinephelus coioides* and role of Sp1-binding site in determining promoter activity. Sci Rep, 8: 5305. DOI 10.1038/s41598-018-23668-w

You, X., Bian, C., Zan, Q., Xu, X., Liu, X., Chen, J., Wang, J., Qiu, Y., Li, W., Zhang, X., Sun, Y., Chen, S., Hong, W., Li, Y., Cheng, S., Fan, G., Shi, C., Liang, J., Tom Tang, Y., Yang, C., Ruan, Z., Bai, J., Peng, C., Mu, Q., Lu, J., Fan, M., Yang, S., Huang, Z., Jiang, X., Fang, X., Zhang, G., Zhang, Y., Polgar, G., Yu, H., Li, J., Liu, Z., Zhang, G., Ravi, V., Coon, S.L., Wang, J., Yang, H., Venkatesh, B., Wang, J. & Shi, Q. (2014). Mudskipper genomes provide insights into the terrestrial adaptation of amphibious fishes. Nature Communications, 5: 5594. DOI 10.1038/ncomms6594

Zheng, X., Ding, Z., Xu, Y., Monroig, O., Morais, S. & Tocher, D.R. (2009a). Physiological roles of fatty acyl desaturases and elongases in marine fish: Characterisation of cDNAs of fatty acyl Δ6 desaturase and elovl5 elongase of cobia (*Rachycentron canadum*). Aquaculture, 290: 122–131.

Zheng, X., Leaver, M.J. & Tocher, D.R. (2009b). Long-chain polyunsaturated fatty acid synthesis in fish: Comparative analysis of Atlantic salmon (*Salmo salar* L.) and Atlantic cod (*Gadus morhua* L.) Delta6 fatty acyl desaturase gene promoters. Comp Biochem Phys B, 154: 255–63. DOI 10.1016/j.cbpb.2009.06.010

Zheng, X., Seiliez, I., Hastings, N., Tocher, D.R., Panserat, S., Dickson, C.A., Bergot, P. & Teale, A.J. (2004). Characterization and comparison of fatty acyl Δ 6 desaturase cDNAs from freshwater and marine teleost fish species. Comp Biochem Phys B, 139: 269–279.

