## Supplementary for "Functional characterization of fatty acyl desaturase Fads2 and Elovl5 elongase in the Boddart’s goggle-eyed goby, *Boleophthalmus boddarti* (Gobiidae) suggest an incapacity for long-chain polyunsaturated fatty acid biosynthesis"

```
Elovl2Cmilii          LGTKFMKDRPAFFLRTHLIIYNLGVMLLS		94
Elovl2Spilchardus     LGPVYMKHRPAYNLKSVLVVYNFSVTMLS		89
Elovl2Omykiss         LGSKYMRNRPAYSLKGVLQVYNFSVTMLS		89
Elovl2Bgonionotus     LGTKYMRNRPAYSLKNILLLYNFSITMLS		89
Elovl2Drerio          LGTKYMRNRPAYSLKNVLLLYNFSVTVLS		89
Elovl2Cgariepinus     IGPKYMKDKPAYSLKNVLLLYNFGVTMLS		94
Elovl2Amexicanus      LGPKYMKNRPAYSLKNVLQLYNFVVTMLS		92
Elovl2Comacropomum    LGPKYMKNRPAYSLKNVLLLYNFSVTMLS		94
Elovl2Loculatus       FGTLYMKNRPAYSLKGVLIAYNFAVTMLS		89
Elovl2Ajaponica       LGPIYMKNRPAYSLKKLLLVYNFAVTMLS		89
Elovl2Sformosus       LGPLYMKNRPAYSLKNVLLLYNFSVTMLS		89
Elovl5Bboddarti       MGPKYMKNRPPFSCRGLLLIYNVGLTLLS		87
Elovl5Scanalicu       LGPKYMKNRPAYSCRGLMVIYNLGLTLLS		86
Elovl2/5Blanceolatum  LGPKLMRSRQAFSLRKVMILYNIIIAALS		86
Elovl2Pmarinus        LGSACMRNRQPLSLRASMVVYNFLVTLLS		86
Elovl5Pmarinus        VGPKLMRERQPFSLKGLLVVYNALLTALS		91
Elovl5Omykiss         LGPKYMRHRQPVSCQGLLVLYNLALTLLS		86
Elovl5Bgonionotus     MGPKYMKNRQPYSCRALLVPYNLCLTLLS		86
Elovl5Danio           MGPKYMKNRQAYSCRALLVPYNLCLTLLS		86
Elovl5Amexicanus      MGPKYMNNRQPFSCRRILVVYNLALTLLS		86
Elovl5Cmacropomum     MGPKYMKDRQPYSCRRILVVYNLALTLLS		86
Elovl5Cgariepinus     MGPKYMRNRQPFSCRGILVLYNLALTFLS		86
Elovl5Sformosus       MGPKYMRSRQAFSCRGLLVIYNLSLTLLS		87
Elovl5Spilchardus     MGPRYMKNRQPISCRGLLVVYNLGLTLLS		86
Elovl5Ajaponica       MGPKYMKNRQPFSCRGLLVVYNLGLTLLS		86
Elovl5Loculatus       IGPKYMNNRQPFSCRGILVIYNLGLTLLS		86
Eovl5Sargus           MGPKYMKYRQPYSCRGLLVFYNLGLTLLS		86
Elovl5Cmilii          AGPKYMKNKAPVSCRGTLVVYNIGLTLLS		86
```

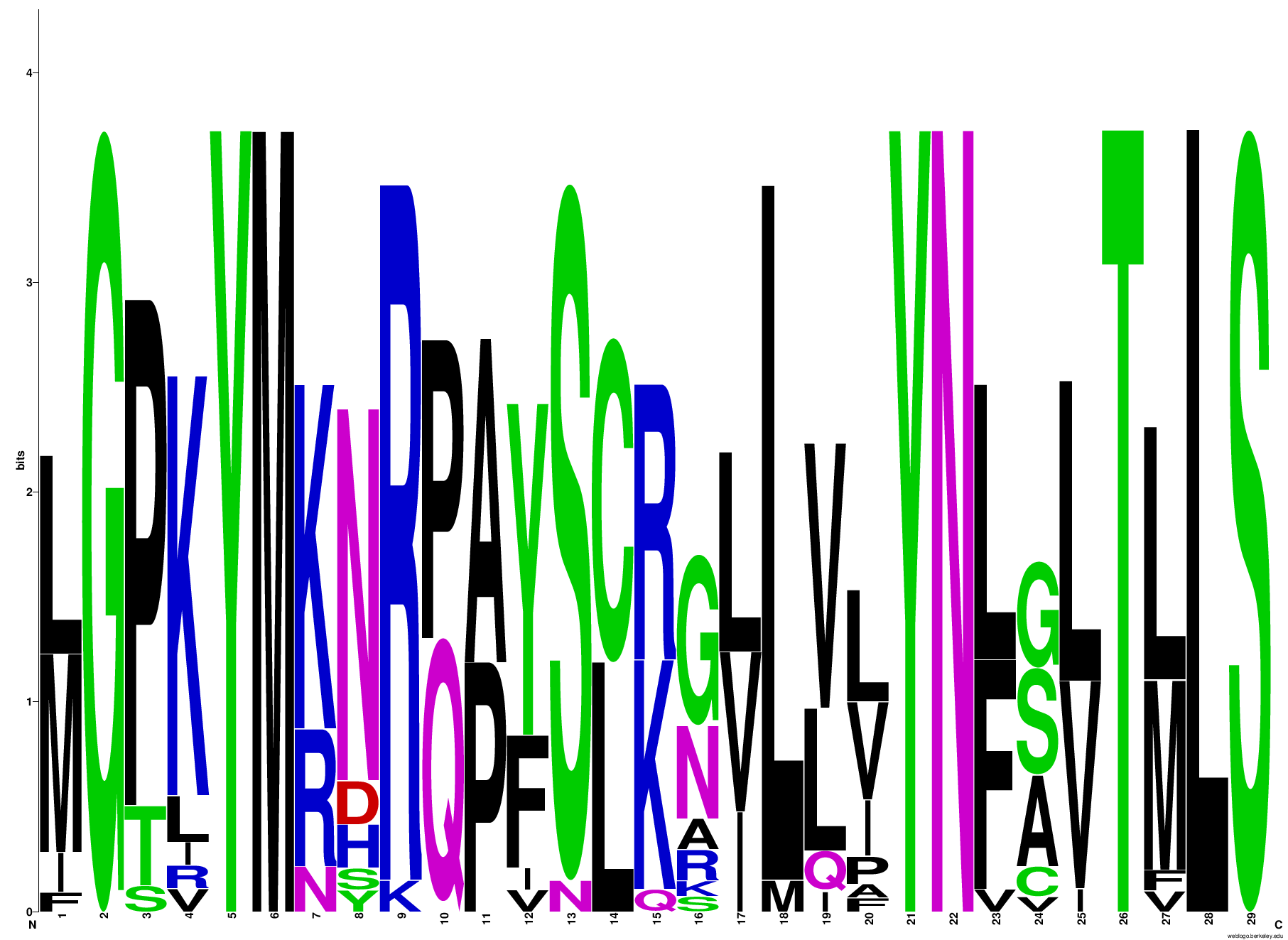

**Supplement 1**

Clustal alignment of partial Elovl5 and Elovl2 amino acid sequences from various species*. Boleophthalmus boddarti* sequence is highlighted in red. Residues corresponding to mutation study are highlighted in blue. The sequence logo was generated with 10 teleost Elovl2 and Elovl5 amino acid sequences. The height of each stack indicates the sequence conservation at that position (measured in bits), whereas the height of symbols within the stack reflects the relative frequency of the corresponding amino acid at that position (Crooks et al 2004). Sequence logos were generated using Weblogo, version 2.8.2., (http://weblogo.berkeley. edu/logo.cgi). Crooks, G.E., Hon, G., Chandonia, J.M. & Brenner, S.E. WebLogo: a sequence logo generator. *Genome Res.* **14**, 1188–1190 (2004).
